# Kindlin2-mediated phase separation underlies integrin adhesion formation

**DOI:** 10.1101/2020.07.10.197400

**Authors:** Yujie Li, Ting Zhang, Huadong Li, Haibin Yang, Ruihong Lin, Kang Sun, Lei Wang, Jing Zhang, Zhiyi Wei, Cong Yu

**Affiliations:** Department of Biology, Southern University of Science and Technology, Shenzhen, Guangdong, China; Academy for Advanced Interdisciplinary Studies, Southern University of Science and Technology, Shenzhen, Guangdong, China; Faculty of Health Sciences, University of Macau, Macau SAR 999078, China; Guangdong Provincial Key Laboratory of Cell Microenvironment and Disease Research, and Shenzhen Key Laboratory of Cell Microenvironment, Shenzhen, Guangdong, China

## Abstract

Formation of cell-extracellular matrix adhesion requires assembly of the transmembrane receptor integrins and their intracellular activators, kindlin and talin proteins in minutes. The mechanisms governing the rapid formation and dynamics of the adhesion remain enigmatic. Here, we reported that the dimerized-kindlin2 underwent phase separation with clustered-integrin in solution and on lipid bilayer. The kindlin2/integrin condensate can further enrich other components for the adhesion complex assembly. The full-length structure of kindlin2 was solved and revealed that the kindlin2 dimers can further pack with each other to form a higher oligomer. Disrupting the intermolecular interaction between the kindlin2 dimer inhibits the phase formation on 2D membrane in vitro and impaired the adhesion formation, integrin activation, and cell spreading in cultured cells. We also determined the full-length structure of kindlin2 in its monomeric conformation. Structural analysis and biochemical characterization indicate that the interdomain interaction control the monomer-dimer transition of kindlin2, providing a regulation mechanism of the kindlin2-mediated phase separation. Our findings not only provide a mechanistic explanation for the formation and dynamic regulation of the integrin-based adhesion, but also shed light on understanding of how the clustered receptors participate in assembly of the functional membrane domains via phase separation.

## Introduction

Cell-extracellular matrix (ECM) adhesions are dynamic subcellular structures connecting ECM and cytoskeletons and mediating the inside-out and outside-in bidirectional communications^1,2^. As the major mediator of cell-ECM adhesions, the transmembrane receptor integrins, comprised of α and β subunits, play critical roles in development, immune response, and hemostasis^3-5^. To initiate the adhesion formation, the integrin receptors are activated and clustered, resulting in the recruitment of over two hundreds of intracellular proteins to the integrin cluster and form a highly dense protein assembly^6,7^. The nascent adhesion or focal complex then either disappears quickly in minutes or eventually matures into the focal adhesion (FA) within half an hour to form a tight connection with the ECM^8,9^. Hundreds of proteins in the focal complex are tightly organized during cell adhesion and migration^10-12^. However, the molecular mechanism underlying the formation and dynamic regulation of the integrin-based adhesion remains elusive.

Identified as integrin-binding proteins, kindlin family proteins play important roles in integrin activation and clustering^13-18^. All kindlins contain a FERM domain, comprised of the canonical F1/2/3 lobes plus an N-terminal F0 lobe, and a unique PH domain inserted in the F2 lobe of the FERM domain (Fig. 1a). The F3 lobe interacts directly to the cytoplasmic tail of β-integrin. In addition to the F3 lobe, the PH domain is also important for localizing kindlins to FA by binding to phosphatidylinositol 3,4,5-trisphosphate (PIP_3_), which is enriched at the FA region^19,20^. Mammalian kindlins have three members, kindlin1/2/3. Among them, kindlin2 is ubiquitously expressed, while kindlin1 and kindlin3 is mainly expressed in epithelia and hematopoietic system, respectively^21,22^. Dysfunction mutations in kindlins associate with many human genetic diseases (e.g. immunity defects, skin defects, and kidney defects) and cancers^23-27^. As one of the earliest arrived components in the integrin-based adhesion^9^, kindlin2 plays a central role in recruiting other FA proteins, such as paxillin and ILK (<u>I</u>ntegrin-linked <u>k</u>inase)^28-30^. Disruption of the kindlin2-binding to β-integrin causes severe defects in the adhesion formation^14,31^, despite the weak association between kindlin2 and β-integrin with a dissociation constant around tens of μM^32^. It’s intriguing to know how such a weak interaction determines the rapid and stable assembly of the integrin-based adhesion.

**Fig. 1.**
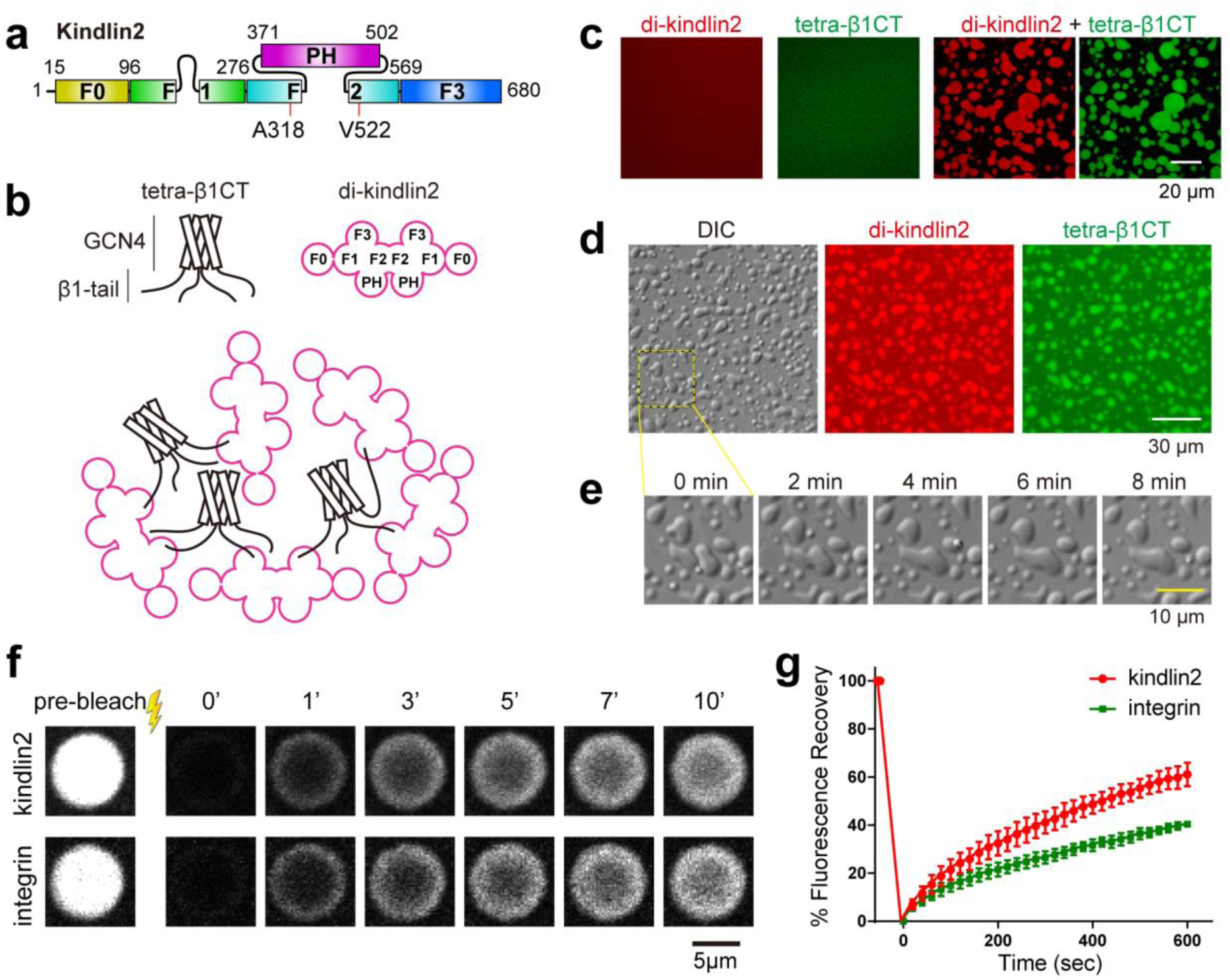
Clustered integrin and dimerized kindlin condensates through phase separation. **a**, Domain organization of kindlin2. **b**, Schematic diagram of the disulfide-linked kindlin2 dimer and the GCN4-tetramerized β1-integrin tail in forming phase separation via the multivalent binding. **c**, Confocal microscopy showing 20 μM cy3-di-kindlin2 mixed with 20 μM Alexa 488-tetra-β1CT formed droplets in solution (right) while neither protein alone at 80 μM could (left). The protein concentration is defined as the unlabeled monomeric unit concentration of each protein throughout the entire study. The labeling ratio is ∼2%. **d**, Differential interference contrast microscopy (left) and widefield fluorescence microscopy (middle and right) of liquid droplets formed by di-kindlin2 and tetra-β1CT at 30 μM with 10 mM DTT in a homemade chamber. **e**, Zoom-in view of the boxed region in panel **d** showing time dependent fusion events of liquid droplets. **f**, FRAP analysis of the kindlin2/integrin droplets. **g**, Quantitative analysis of the fluorescence signal shown in panel **f**. Error bars were obtained by analyzing 6 droplets.

Recently, we reported that kindlin2 can adopt a dimer conformation via the F2 lobe to activate integrin and the monomer-dimer transition of kindlin2 plays a critical role in the integrin-based adhesion^32^. In this study, we discovered that the dimerized-kindlin2 forms condensates with the clustered β-integrin via phase separation in solution and on reconstituted 2D membrane. We further found that the PH domain is essential for the phase separation on the 2D membrane system. By solving the crystal structure of the full-length kindlin2 protein, we identified the intermolecular packing between the PH domain and F0 lobe, which induces the dimerized-kindlin2 to form a higher oligomer and promotes the phase separation of β-integrin and kindlin2 on the 2D membrane. The phase formation allows other FA proteins to enter the integrin-dense condensates. Cellular analysis showed that the kindlin2 condensate formation is required for the integrin-based adhesion formation and dynamics as well as integrin-mediated cellular processes. Finally, structural comparison of the full-length kindlin2 structures in the monomer and dimer conformations suggests that the formation of the kindlin2 condensate depends on the monomer-dimer transition of kindlin2. Taken together, our results demonstrated that the phase separation mediated by the clustered-integrin and oligomerized-kindlin2 orchestrates the FA components for the integrin adhesion formation.

## Results

### Phase separation of kindlin2 dimer and clustered β1-integrin in solution

Our previous study uncovered that kindlin2 promotes the activation of clustered integrins through forming a domain-swapped dimer^32^. We then tried to test whether the dimerization of kindlin2 could enhance the binding of kindlin2 to the cytoplasmic tail of the clustered β1-integrin. To mimic the clustered integrin tail, we fused the cytoplasmic tail of β1-integrin (residues 756-798) to the coiled-coil domain of GCN4 (named tetra-β1CT) ^33^, which forms a stable tetramer (Extended Data Fig. 1a). The dimer formation of kindlin2 took weeks in solution^32^. To accelerate the process, we generated the modified dimer (named di-kindlin2) by employing the designed disulfide-crosslink in the dimerization interface (Extended Data Fig. 1b&c). Unexpectedly, when mixing tetra-β1CT and di-kindlin2 at a protein concentration of 30 μM, we found the formation of opalescent solution (Extended Data Fig. 1d) that prevents us from measuring the binding affinity. As several types of liquid-liquid phase separation have been reported to be mediated by multivalent protein-protein interaction^34,35^, the observed opalescent solution likely contains protein condensates formed by phase separation, considering the multi-site binding between tetra-β1CT and di-kindlin2 (Fig. 1b).

To investigate the potential phase formation, we mixed fluorescently labeled tetra-β1CT (Alexa Fluor 488) and di-kindlin2 (Cy3) at a 1:1 ratio and analyzed the mixture using light microscopy. Indeed, numerous spherical-shaped and micron-sized droplets were observed either in the 96 plate or in the home-made chambers, as shown by confocal, differential interference contrast (DIC), and wide field fluorescence images (Fig. 1c&d). Both tetra-β1CT and di-kindlin2 in the mixture were highly enriched in the droplets, while no such droplets could be observed in tetra-β1CT or di-kindlin2 alone (Fig. 1c). Time-lapse imaging further showed that the droplets can fuse with each other (Fig. 1e). Fluorescence recovery after photobleaching (FRAP) assay also revealed that tetra-β1CT or di-kindlin2 in the droplets was constantly exchanging with protein molecules in the surrounding dilute solution (Fig. 1f&g). Thus, the microscopic analysis confirmed that tetra-β1CT and di-kindlin2 underwent phase separation in solution.

### The phase formation depends on the kindlin2/β1-tail interaction as well as the oligomerization of kindlin2 and β1-tail

As neither tetra-β1CT nor di-kindlin2 alone undergoes phase separation, the observed phase formation likely requires the complex formation between kindlin2 and β1-tail. Consistently, no phase formation was observed for tetra-β1CT carrying the kindlin2-binding deficient mutations (T788A and Y795A) when mixing with di-kindlin2 (Fig. 2a and Extended Data Fig. 2).

**Fig. 2.**
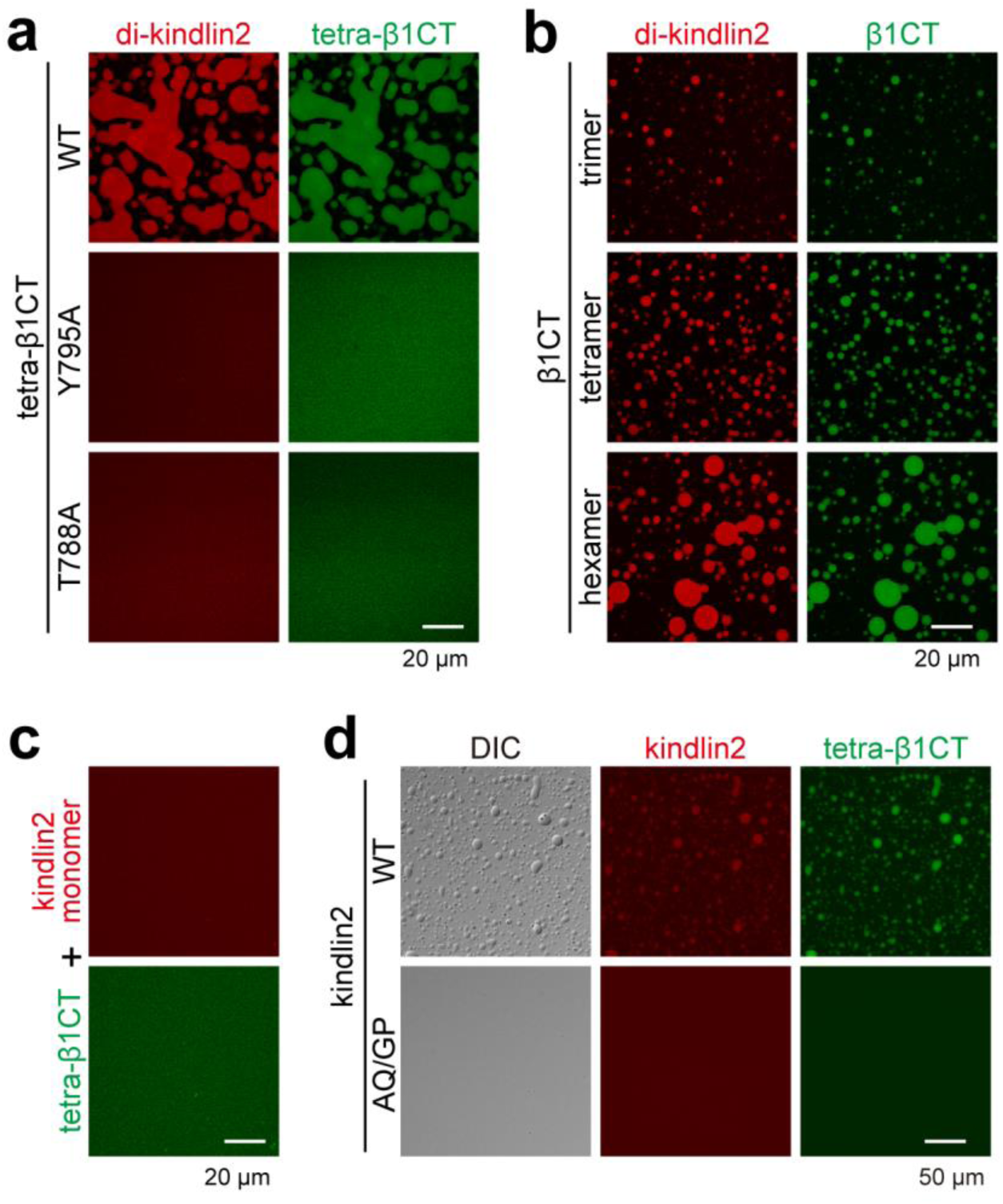
Phase formation depends on the kindlin2/β1-tail interaction and the oligomerization state of kindlin2 and β1-tail. **a**, Confocal microscopy of phase separation prepared by mixing di-kindlin2 with either tetra-β1CT wild-type or its kindlin2-binding deficient mutations (Y795A and T788A) at the concentration of 20 μM. **b**, Confocal microscopy of phase separation formed by di-kindlin2 and β1CT with different oligomerization states at 10 μM each. **c**, Monomeric kindlin2 cannot undergo phase separation with tetra-β1CT at 80 μM each. **d**, The AQ/GP mutation on kindlin2 interferes with the phase separation as observed by DIC and fluorescence imaging. As the AQ/GP mutation inhibits the disulfide-linked kindlin2 dimer, the samples were prepared by direct adding the 40 μM mixture of the A318C and V522C mutants of kindlin2 with or without the AQ/GP mutation into tetra-β1CT.

Adding di-kindlin2 to the trimer, tetramer or hexamer of β1-tail, the number and size of the protein condensates increased with the increased oligomerization states of β1-tail (Fig. 2b), indicating that the condensate formation depends on the oligomerization of β1-tail. Likewise, the dimer formation of kindlin2 is required for the phase separation. Mixing of the purified kindlin2 monomer with tetra-β1CT failed to generate the condensates even at the protein concentration of 80 μM (Fig. 2c). Consistently, the AQ/GP mutation in kindlin2 that was found to disrupt the monomer-to-dimer transition of kindlin2^32^ diminished the phase separation phenomenon of kindlin2 with tetra-β1CT (Fig. 2d).

### Di-kindlin2 forms condensates with tetra-β1CT on lipid membrane bilayer

In living cells, the clustered integrin receptors recruit a large number of cytoplasmic proteins and serve as a membrane-associated platform for signal transduction. To understand the role of the observed phase separation for such a membrane-attached assembly, we reconstituted the tetra-β1CT on a 2D membrane system using supported lipid bilayers with the defined lipid composition (Fig. 3a)^36,37^, and analyzed the phase separation formed by tetra-β1CT and di-kindlin2 in the membrane environment.

**Fig. 3.**
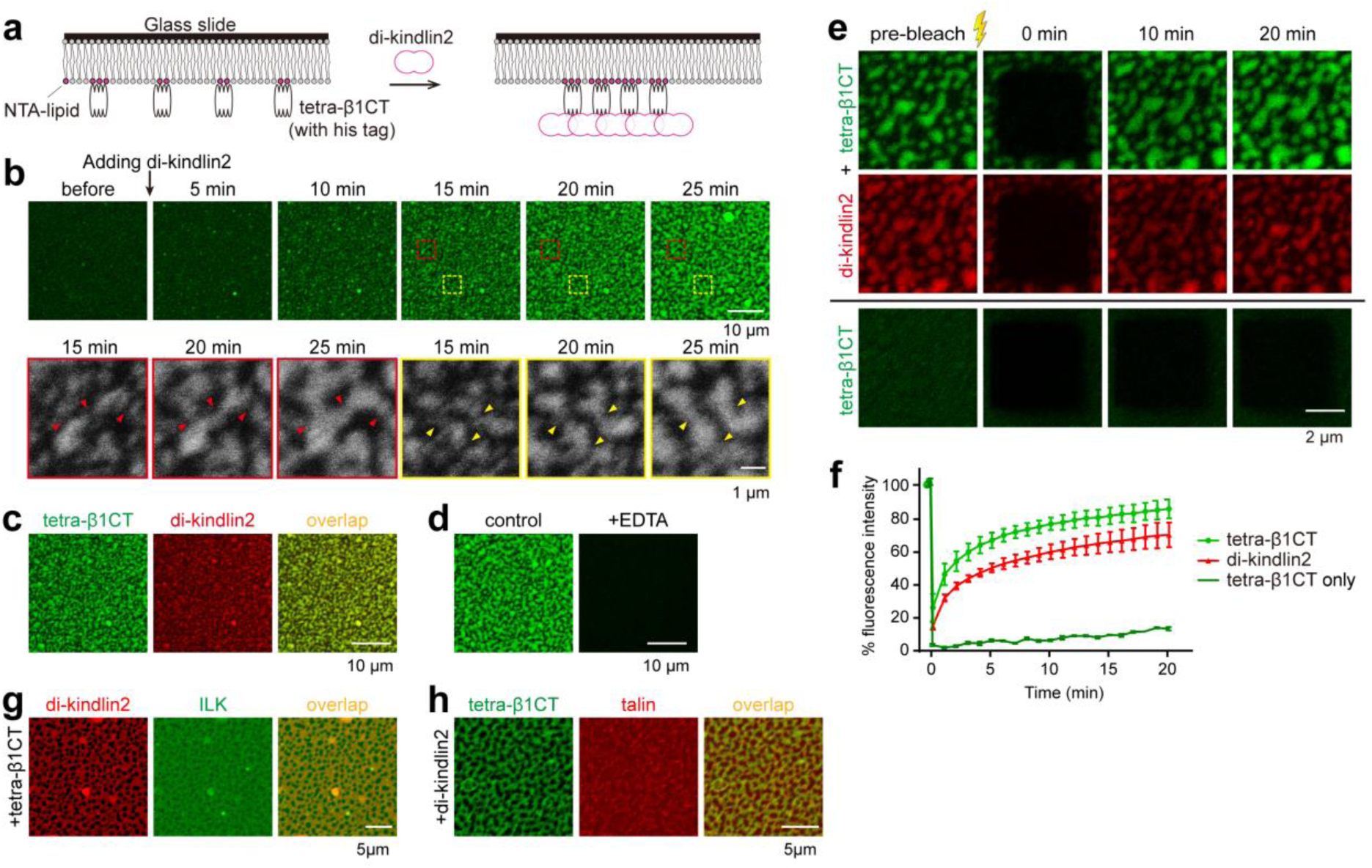
Reconstitution of kindlin2/integrin condensates on the supported lipid bilayer. **a**, Cartoon diagram of the experimental design. The His-tagged tetra-β1CT was attached to lipid bilayer on coverslip by NTA lipid and di-kindlin2 was subsequently added to the system. **b**, Time dependent imaging of His-tetra-β1CT on supported lipid bilayer (SLB) after addition of di-kindlin2 to the final concentration of 2 μM. Two selected regions were enlarged. Arrowhead indicate the condensates fused into network. Images were taken by confocal microscopy. **c**, Confocal microscopic analysis of condensates on SLBs containing both His-tetra-β1CT and di-kindlin2. The images were taken at 25 minutes after adding di-kindlin2 to His-tetra-β1-attached SLB. **d**, Addition of 2mM EDTA to the system abolishes the phase formation on SLB. **e**, FRAP analysis of His-tetra-β1CT and di-kindlin2 formed condensates on SLB. His-tetra-β1CT alone was used as control. **f**, Quantification of panel **e. g&h**, His-tetra-β1CT and di-kindlin2 on SLB enriched the ILK/Parvin complex **g** and talin_head **h** into the condensates.

The tetra-β1CT protein captured by the Ni^2+^-NTA-DGS-containing lipid bilayers was uniformly distributed (Fig. 3b). Upon adding di-kindlin2, condensates appeared within a few minutes. Small condensates with enriched tetra-β1CT and di-kindlin2 gradually grew larger and became irregular shapes on the membrane surface. Finally, the condensates coalesced into a mesh-like network (Fig. 3b&c). The dissociation of tetra-β1CT from membrane by adding 2 mM EDTA together with di-kindlin2 in the system abolished condensate formation (Fig. 3d), indicating that the membrane-association of tetra-β1CT is required for the phase formation. Similar with our observation of the phase separation in solution, both the fluorescently labeled tetra-β1CT and di-kindlin2 in the membrane condensates exhibited dynamic exchanges with those in the membrane or aqueous solution, respectively, as indicated by the high recovery rate after photobleaching in FRAP analysis (Fig. 3e&f). In contrast, little recovery was observed when bleaching the membrane-bound tetra-β1CT only (Fig. 3e&f). These results reveal that the complex of tetra-β1CT and di-kindlin2 have the capability of phase separation on lipid membrane bilayer as well.

### PH domain plays a unique role in the kindlin2-mediated phase separation

Compared with other FERM-containing proteins, kindlins contain a unique PH domain, inserted in the F2 lobe (Fig. 1a). Although the conformational change of the F2 lobe is required for the kindlin2 dimerization, the deletion of the PH domain does not compromise the dimer formation of kindlin2 (Extended Data Fig. 1e)^32^. Surprisingly, removing the PH domain from di-kindlin2 abolishes the formation of the irregular-shaped clusters and the mesh-like network on the 2D membrane, although a few round-shaped droplets were observed near the membrane layer by using confocal microscope (Extended Data Fig. 3a). Our observations suggest that the PH domain is essential for the cluster formation on membrane but not required for the phase separation in solution. Consistently, the condensate formation in solution was not impaired by the deletion of the PH domain in kindlin2 (Extended Data Fig. 3b-d). Although the PH domain has been implicated to be involved in the membrane binding via the interaction with phosphatidylinositol 3,4,5-trisphosphate (PIP_3_) or phosphatidylinositol 4,5-trisphosphate (PIP_2_)^19,20^, it is unlikely that the PH domain controls the phase formation on the membrane through the interaction with PIP_3_ or PIP_2_, as no PIPs was included in our lipid bilayer preparation.

### The intermolecular interaction between the PH domain and the F0 lobe is critical for phase separation formation

As the PH domain was deleted in the previously determined kindlin2ΔPH structures, to understand the role of PH domain in the phase separation on the membrane, we solved the full-length structure of kindlin2 (Table 1). Same as the dimeric kindlin2ΔPH structure, the full-length kindlin2 dimerizes in the F2-swapped manner (Fig. 4a and Extended Data Fig. 4a). Although the connecting loops between the PH domain and F2 lobe are largely unstructured, the PH domain in the full-length kindlin2 structure is well-defined and interacts with the F2 lobe intramolecularly (Fig. 4a).

**Table 1.** Data collection and refinement statistics.

|  | **Kindlin2-dimer/IP<sub>6</sub>**<br>**(PDB 6LWW)** | **Kindlin2-monomer**<br>**(PDB 6LWX)** |
| --- | --- | --- |
| **Data collection** |  |  |
| Space group | *C* 2 2 2<sub>1</sub> | *P* 2<sub>1</sub> 2<sub>1</sub> 2<sub>1</sub> |
| Cell dimensions |  |  |
| *a*, *b*, *c* (Å) | 88.0, 124.4, 370.1 | 123.8, 127.0, 175.9 |
| $\alpha$, $\beta$, $\gamma$ (°) | 90, 90, 90 | 90, 90, 90 |
| Resolution (Å) | 50–3.35 (3.41–3.35) | 50–3.20 (3.26–3.20) |
| *R*<sub>merge</sub><sup>a</sup> | 0.104 (1.376) | 0.182 (2.98) |
| *I*/σ*I* | 25.4 (2.2) | 23.5 (1.2) |
| *CC*<sub>1/2</sub><sup>b</sup> | 0.997 (0.601) | 0.996 (0.643) |
| Completeness (%) | 97.9 (99.9) | 97.4 (84.7) |
| Redundancy | 12.1 (12.7) | 27.3 (26.8) |
| **Refinement** |  |  |
| Resolution (Å) | 50–3.35 (3.47–3.35) | 50–3.20 (3.28–3.20) |
| No. reflections | 28971 (2807) | 45020 (2738) |
| *R*<sub>work</sub> / *R*<sub>free</sub><sup>c</sup> | 0.239 (0.369) / 0.298 (0.422) | 0.218 (0.356) / 0.260 (0.363) |
| No. atoms | 8678 | 9357 |
| Protein | 8622 | 9341 |
| Ligand/ion | 56 | 10 |
| Water | 0 | 6 |
| Mean *B* (Å<sup>2</sup>) | 177.0 | 131.1 |
| Protein | 176.5 | 131.1 |
| Ligand/ion | 248.3 | 150.2 |
| Water | - | 90.0 |
| r.m.s. deviations |  |  |
| Bond lengths (Å) | 0.003 | 0.002 |
| Bond angles (°) | 0.84 | 0.61 |
| Ramachandran analysis |  |  |
| Favored region (%) | 96.4 | 95.2 |
| Allowed region (%) | 3.6 | 4.8 |
| Outliers (%) | 0 | 0 |

The numbers in parentheses represent values for the highest resolution shell.
<sup>a</sup>*R*<sub>merge</sub> = $\sum |I_i - I_m| / \sum I_i$ , where *I*<sub>i</sub> is the intensity of the measured reflection and *I*<sub>m</sub> is the mean intensity of all symmetry related reflections.
<sup>b</sup>*CC*<sub>1/2</sub> is the correlation coefficient of the half datasets.
<sup>c</sup>*R*<sub>work</sub> = $\sum ||F_{obs}| - |F_{calc}|| / \sum |F_{obs}|$ , where *F*<sub>obs</sub> and *F*<sub>calc</sub> are observed and calculated structure factors.
*R*<sub>free</sub> = $\sum_T ||F_{obs}| - |F_{calc}|| / \sum_T |F_{obs}|$ , where *T* is a test data set of about 4–5 % of the total reflections randomly chosen and set aside prior to refinement.

**Fig. 4.**
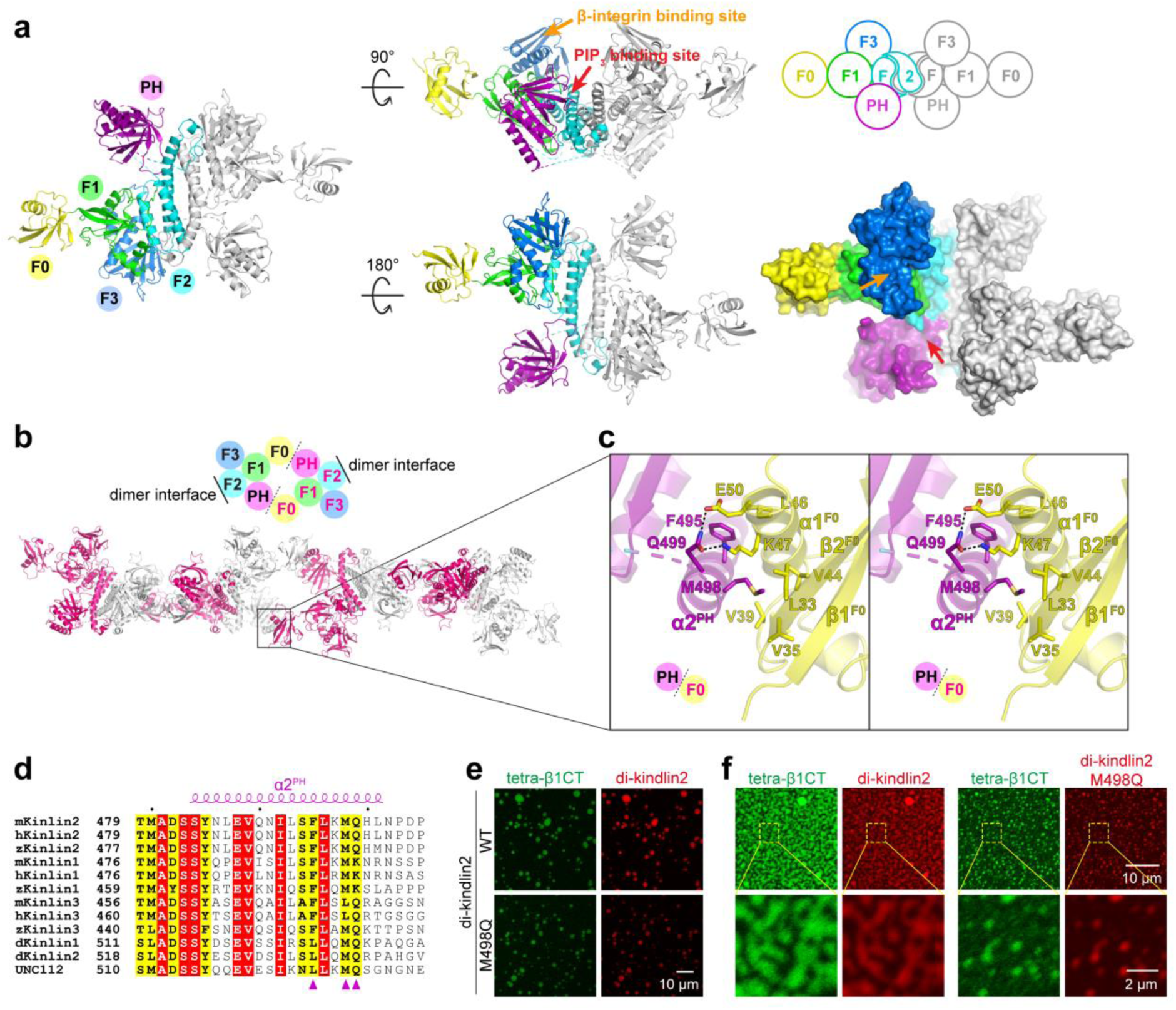
Structural basis of the PH domain-mediated kindlin2 oligomerization. **a**, Ribbon representations of the kindlin2 dimer structure with three views. One protomer is colored following the coding scheme in Fig. 1a and the other identical protomer is colored in gray. The disordered loops in F1 and in the connecting region between PH and F2 is indicated by dashed lines. The cartoon of kindlin2 dimer schematically shows the interdomain interactions and the F2 lobe-swapped dimer. The surface analysis of kindlin2 indicates the deeply positioned PIP_3_-binding pocket in the PH domain. **b**, The kindlin2 polymer in crystal is formed through the intermolecular PH/F0 interaction between the kindlin2 dimers. **c**, Stereoview of molecular details in the PH/F0 interface. Hydrogen bonds and salt bridges were indicated by dotted lines. **d**, Multiple sequence alignment of the C-terminal helix in the PH domain of kindlin2 from different species. The species were represented by “h” for human, “m” for mouse, “z” for *Danio rerio*, “d” for *Drosophila melanogaster*. UNC112 is the kindlin2 homologue in *Caenorhabditis elegans*. Residues involved in the interaction with the F0 lobe are indicated by solid triangles. **e**, Phase separation of di-kindlin2 WT or M498Q and tetra-β1CT in solution. Di-kindlin2 and tetra-β1CT were mixed at 20 μM each, and then diluted 4 times with the same buffer. **f**, Condensates on SLB at 25 minutes after adding 2 μM di-kindlin2 WT or M498Q to His-tetra-β1CT attached SLB.

Interestingly, crystal packing analysis showed that the PH domain packs with the F0 lobe by intermolecular interactions (Fig. 4b). The PH/F0 packing is mainly mediated by hydrophobic interactions, in which M498^PH^ in the C-terminal α-helix (α2^PH^) of the PH domain inserts its sidechain into a hydrophobic pocket formed by several hydrophobic residues in the F0 lobe (Fig. 4c&d). In addition, Q499^PH^ facilitates the intermolecular packing by forming two hydrogen bonds with E50^F0^ and K47^F0^ (Fig. 4c) and drags the sidechain of K47^F0^ away from the hydrophobic pocket of the F0 lobe observed in the kindlin2ΔPH structures, to avoid the potential clash between K47^F0^ and M498^PH^ (Extended Data Fig. 4b). The PH/F0 packing and the F2-swapped dimerization together results in the formation of the kindlin2 polymer in crystal (Fig. 4b). Consistently, although the PH/F0 interaction is too weak to be detectable in solution (Extended Data Fig. 4c), the di-kindlin2 protein further oligomerized after being placed at 4 °C for two days, whereas the oligomer fraction of di-kindlin2ΔPH was hardly detectable even after 9 days (Extended Data Fig. 4d&e). Given the importance of the multivalent interaction in phase separation^34,38-40^ and the involvement of the PH domain in the phase formation on the membrane, the additional PH/F0 intermolecular interaction is likely to play a role in the kindlin2-mediated phase formation on the membrane.

To test whether the PH/F0 packing contributes to the phase formation, we designed a mutation at the F0-binding surface of the PH domain by substituting M498 to glutamine (M498Q), which presumably disrupts the hydrophobic interaction between the PH domain and the F0 lobe. Di-kindlin2 with the M498Q mutation retains the capability to form the phase with tetra-β1CT in solution (Fig. 4e and Extended Data Fig. 1f) and moderately decreased the size of the droplets (Extended Data Fig. 3c&d). However, mesh-like network of phase on the 2D membrane system was largely attenuated by using the M498Q mutant (Fig. 4f). Thus, the intermolecular interaction between di-kindlin2 molecules through the PH domain and the F0 lobe likely promotes the phase formation by increasing the binding valency, especially in the lipid bilayer environment.

### Di-kindlin2 and tetra-β1CT form the condensates together with other FA components

Since β-integrin and kindlin2 are capable of binding to a number of FA proteins, including talin and ILK, respectively^28,41,42^, it is likely that talin and ILK can be incorporated into the condensates formed by the clustered β-integrin and the kindlin2 dimer. The ILK fragment containing the kinase domain was co-purified with the CH2 domain of parvin^43^. The binding of the ILK/parvin complex to kindlin2 was confirmed (Extended Data Fig. 5a). Upon mixing tetra-β1CT and di-kindlin2, the ILK/parvin complex were concentrated in the condensed droplets (Extended Data Fig. 5b). Likewise, as the N-terminal FERM domain of talin (talin head) directly interacts with β1-tail, the talin head fragment was concentrated into the phase formed by tetra-β1CT and di-kindlin2 (Extended Data Fig. 5c). Similarly, the ILK/parvin complex and talin head could also be enriched in the condensates formed by di-kindlin2 and tetra-β1CT on the lipid membrane bilayer (Fig. 3g&h). These results indicate that the condensed phase formed by kindlin and integrin is capable of enriching other FA proteins, such as talin and ILK.

### Kindlin2-mediated phase separation is critical for the formation and dynamics of the integrin-based adhesion

As the formation of the integrin-based adhesion are minute-scale processes, the kindlin2-mediated phase separation observed *in vitro* provide a potential mechanism to achieve such a rapid assembly in cells. Consistent with many previous studies showing the highly dynamic feature of the nascent adhesion and the mature FA in various types of cells^40,44^, we observed that fusion event occurs during the formation of the FAs in the kindlin2-knockout HT1080 cells transfected with kindlin2 (Extended Data Fig. 6a, Movie S1). FRAP experiments confirmed that kindlin2 molecules at the FA exchanged rapidly with those in the cytoplasm, as observed by others (Extended Data Fig. 6b)^44^. These data together suggested that the kindlin2-mediated phase separation occurs at the integrin-based adhesion.

Our biochemical and structural characterization showed that the oligomerization of kindlin2 is essential for the phase separation of β-integrin tails (Fig. 4). To further investigate the kindlin2-mediated phase separation in cells, we performed a series of cellular assays by using the kindlin2-knockout HT1080 cell transfected with either the wild-type kindlin2 or its two mutants, AQ/GP and M498Q, which show defects in the dimerization and oligomerization of kindlin2, respectively (Fig. 4b)^32^. As M498^PH^ locates at a region distal to the PIP_3_-binding site of the PH domain (Extended Data Fig. 7a), the M498Q mutation is unlikely to interfere with the lipid binding of kindlin2 mediated by the PH domain.

As expected, the AQ/GP and M498Q mutants of kindlin2 localized to the FA as indicated by paxillin staining (Fig. 5a), suggesting that these mutants retain their capability to interact with β-integrin. However, both the size and number of FA in the cells overexpressing the mutants were decreased compared with the cells overexpressing the wild-type kindlin2 (Fig. 5a&b). We further noticed that, after cells were plated on fibronectin coated glass surface, FAs were generated within a few minutes in the cells overexpressing the wild-type kindlin2. In contrast, in the cells that were transfected with the kindlin2 mutants, although the number of the newly formed adhesions remains unchanged, the growth rate of these adhesions was largely decreased, especially in the cells expressing the M498Q mutant, as indicated by the size change of the adhesions during 5 minutes after newly appeared (Fig. 5c&d, Movies S2-4). In addition, the newly formed adhesions were relatively stable in the extending region of the cells transfected with the wild-type kindlin2, whereas the small adhesions were often disappeared in the cells transfected with the M498Q mutant (Extended Data Fig. 6c, Movie S5). Together, these results indicated that the proper formation of the integrin-based adhesion requires the kindlin2-mediated phase separation and the mutations impairing the phase formation and dynamics *in vitro* could also impair the integrin-based adhesion *in vivo*.

**Fig. 5.**
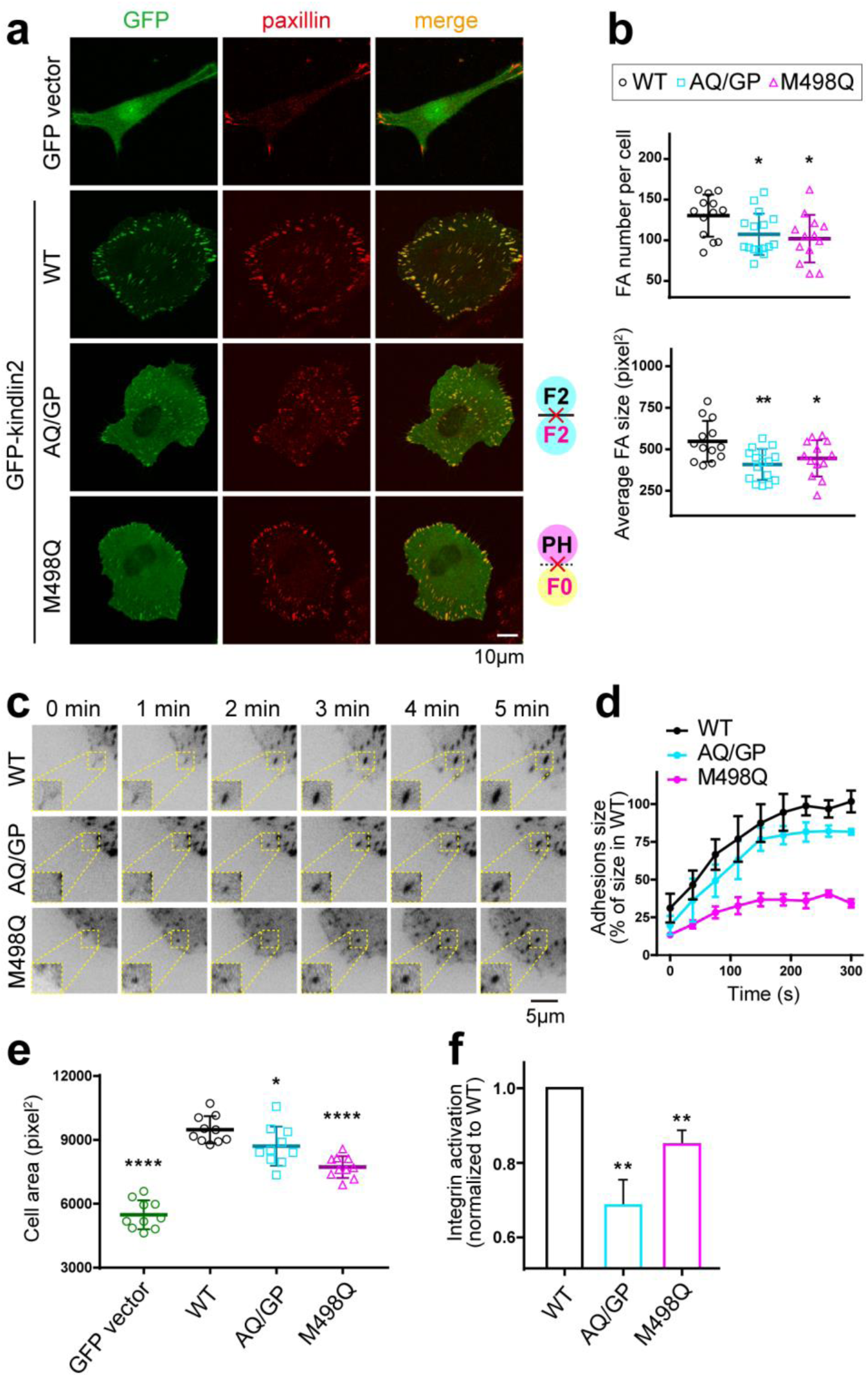
Functional characterization of the kindlin2/integrin condensates in cells. **a**, FA localization analysis of transient-expressed GFP-kindlin2 WT and mutants in kindlin2-KO HT1080 cells. FAs were indicated by paxillin staining. Both the AQ/GP and M498Q mutant were able to localize to FA but reduced FA size and number. **b**, Quantification of FA size and number as shown in panel **A**. FA was counted and measured by paxillin marking 12 to 15 cells per experiment. **c**, TIRF live cell imaging of adhesion formation. The kindlin2-KO cells transfected with GFP-kindlin2 were plated on fibronectin-coated dishes. Images of GFP signal were taken about 15 minutes after plating. Time zero was defined as the starting point of one specific FA. **d**, Quantification of the sizes of the newly formed FA (normalized to the average size of the adhesion in WT) over time was measured. Curve showed average value of about 10 FAs. **e**, Cell spreading analysis of the kindlin2 mutants. Kindlin2-KO cells were transfected with GFP vector or GFP-tagged kindlin2 variants. Effects of kindlin2 mutants on cell spreading were quantified by measuring cell areas. The cell area was calculated as total areas (stained with Phalloidin-555)/number of cells. A two-tailed unpaired Student’s t-test was performed to test for significance between different experimental datasets. Mean value of 10 positions (each position contains 30 ∼ 40 cells) in each sample was calculated. **f**, Integrin activation analysis of the kindlin2 mutants. Individual GFP-tagged kindlin2 WT or variants were co-transfected with RFP-tagged talin-FERM in αIIbβ3-CHO cells. αIIbβ3-integrin activation was evaluated by PAC1 binding. Data were normalized to cells expressing WT kindlin2. All bars represent the means ±S.D.. *, p< 0.05; **, p<0.01; ***, p<0.001; ****, p< 0.0001.

Since the cell spreading highly relies on the adhesion formation, we analyzed the potential cell spreading defects caused by the M498Q or AQ/GP mutations in kindlin2. Indeed, the cell spreading areas were decreased in the cells overexpressing either the M498Q or AQ/GP mutant (Fig. 5e and Extended Data Fig.6d). Consistent with the importance of the oligomerization of kindlin2 in the formation and dynamic regulation of the integrin-based adhesion, neither the M498Q nor AQ/GP mutant was able to fully rescue the kindlin2-mediated co-activation of integrin to the level comparable to the wild-type kindlin2 in the kindlin2 knock-out cells (Fig. 5f). Taken together, the above structural, biochemical and cellular analysis demonstrated that the dimerization/oligomerization of kindlin2 promotes the integrin-based adhesion formation through the phase separation with the clustered integrins.

### The monomeric kindlin2 adopts a conformation for membrane binding

Our above data indicated that the PH domain of kindlin2 involves in the phase separation by mediating the intermolecular interaction between the dimeric kindlin2 molecules (Fig. 4b). Surprisingly, the PIP_3_-binding pocket in the PH domain is deeply positioned in the concave surface of the kindlin2 dimer (Fig. 4a, indicated by a red arrow) and thereby unable to access the head of PIP_3_ in the membrane with little curvature. This structural observation is inconsistent with the role of the PH domain in phospholipid binding^19^. Hence, the plasma membrane recruitment of kindlin2 may require the PH domain to adopt an orientation different from that in the dimer structure.

Our previous structural study of kindlin2ΔPH revealed a large conformational change of the F2 lobe between the monomer and dimer structures of kindlin2ΔPH^32^. As inserted in the middle of the F2 lobe, the PH domain is likely to switch its position between the kindlin2 monomer and dimer. To confirm that the PH domain undergoes a positional change during the monomer-dimer transition, we determined the full-length structure of the kindlin2 monomer in a high salt condition (Table 1). Indeed, the PH domain is reoriented in the monomer structure (Fig. 6a). Interestingly, the regions of kindlin2 reported to be involved in the association with the membrane (e.g. the long loop of the F1 lobe and the PIP_3_-binding pocket of the PH domain^20,45-47^) or the integrin-binding groove of the F3 lobe^32^) are facing to the same plane (Fig. 6b). Thus, such a planar organization of the membrane-association elements in the monomer structure implicates that the kindlin2 monomer adopts the conformation ready for the plasma membrane recruitment (Fig. 6b). We note with interests that the critical PIP_3_-binding loop in the PH domain shows a slightly altered conformation in the full-length structures, compared with those in the individual PH domain structures of kindlins (Extended Data Fig. 7a)^46-48^. Nevertheless, the PH domain in the full-length structures likely possesses the PIP_3_-binding ability, as indicated by the unexpected IP_6_ binding to the PIP_3_-binding pocket (Extended Data Fig. 7b).

**Fig. 6.**
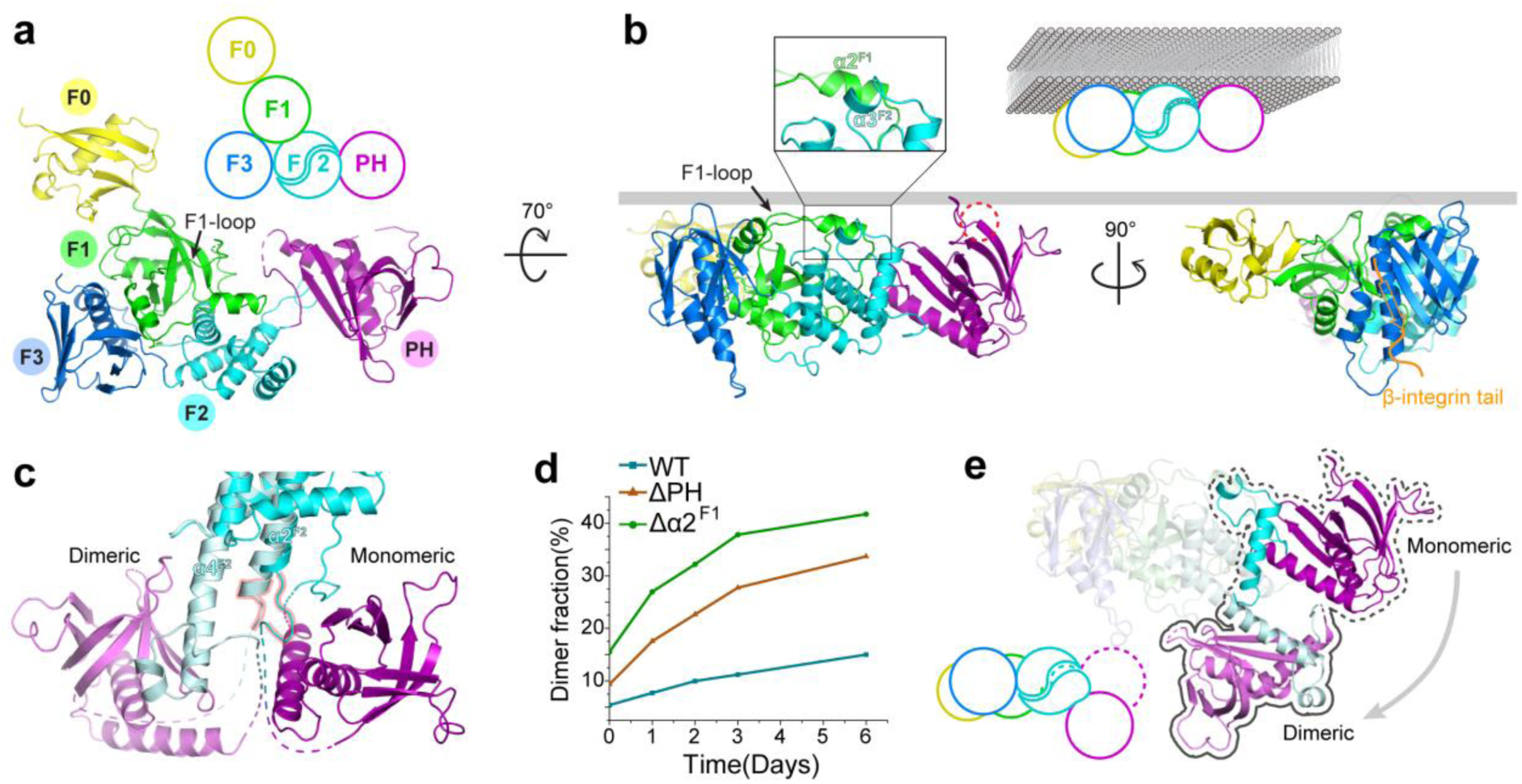
The monomeric structure of kindlin2. **a**, Ribbon representation of the kindlin2 monomeric conformation. **b**, The planar arrangement of the PH domain and the four lobes in the FERM domain. The PIP_3_-binding pocket was highlighted by a dashed circle. The binding site in the F3 lobe for the β-integrin tail was indicated by a modeled β-tail binding, according to the previously determined kindlin2ΔPH/β-tail structure (PDB id: 5XQ0). The membrane binding mode of the kindlin2 monomer was illustrated as a cartoon scheme. The α2^F1^/α3^F2^ interaction was shown as an enlarged view. **c**, Structural comparison of the monomeric and dimeric conformation of kindlin2 showing the distinct interdomain interaction between the F2 lobe and the PH domain. The molecular details of the F2/PH interfaces in the kindin2 monomer and dimer were shown in Extended Data Fig. 9. **d**, Quantification of the dimer fraction of kindlin wild-type and mutations as shown in Extended Data Fig.10. The samples were prepared using the fresh-purified protein as 0 day or the same batch of protein placed at 4 °C for different days. **e**, The large conformational change of the PH domain between the kindlin2 monomer and dimer. The F0, F1, and F3 lobes of the kindlin2 monomer and dimer were superimposed. The region undergoes structural changes were highlighted.

### The intramolecular interactions regulate the monomer-dimer transition of the kindlin2

As the kindlin2-mediated phase formation on the membrane requires the dimer formation of kindlin2, it is essential for kindlin2 to switch from the membrane-associated monomer to dimer. Our previous structural analysis of kindlin2ΔPH showed that the structural transition from the monomer to dimer is caused by the conformational change of the F2 lobe from a compact fold to an extended conformation. However, the kindlin2 monomer is stable in solution and the transition from kindlin2 monomer to dimer is extremely slow *in vitro*^32^. Interestingly, the purified kindlin2 fragment containing the F2 lobe alone predominantly forms a dimer (Extended Data Fig. 8a). It suggests that the dimer formation is an intrinsic property of the F2 lobe and presumably blocked by the interdomain interactions in the full-length kindlin2.

In kindlin2 monomer, the F2 lobe mainly interacts with the PH domain and the F1 lobe (Fig. 6a). The PH domain binds to the F2 lobe in both the monomer and dimer via distinct binding modes (Fig. 6c and Extended Data Fig. 9a&b). In the monomeric conformation, the partially modeled loop region (residues 329 – 334) following the α2^F2^-helix is involved in the intramolecular PH/F2 interaction (Extended Data Fig. 9c). However, the corresponding loop region in the dimeric conformation refolds as the C-terminal part of the α2^F2^-helix, which tightly packs with and stabilizes the long, straight α4^F2^-helix (Fig. 6c and Extended Data Fig. 9d). As forming the α4^F2^-helix is essential for the F2-mediated dimerization of kindlin2, it is likely that the PH domain, by binding to this loop region in the kindlin2 monomer, elevates the energy barrier for the conformational change of the F2 lobe and thereby inhibits the monomer-dimer transition. In line with our hypothesis, the dimerization fraction in the purified kindlin2 fragment containing the F2 lobe with the PH domain insertion was largely decreased (Extended Data Fig. 8b). In the full-length protein, deletion of the PH domain also increased the dimerization rate of kindlin2 (Fig. 6d and Extended Data Fig. 10a&b).

In addition to the PH domain, an inserted long loop (F1-loop) in the F1 lobe is also involved in the intramolecular binding to the F2 region that shows a large structural change between the monomeric and dimeric conformation (Fig. 6b&e). Specifically, an α-helix (α2^F1^) at the C-terminus of the F1 loop packs with a short α-helix (α3^F2^) of the F2 lobe in the kindlin2 monomer. To form the F2-swapping dimer, the α3^F2^-helix has to be intramolecularly uncoupled with the α2^F1^-helix and then intermolecularly recoupled with the α2^F1^-helix again (Fig. 6b&e)^32^. Such an intramolecular coupling between these two helices may inhibit the dimer formation of kindlin2. Consistently, deleting this short α2^F1^-helix (residues 226 – 237) in kindlin2 accelerates the dimer formation (Fig. 6d and Extended Data Fig. 10c). Together, our structural and biochemical analysis indicates that the intramolecular interaction between the F2 lobe and the PH domain or the F1-loop attenuates the conformational change of kindlin2 from monomer to dimer.

## Discussion

Many transmembrane receptors function as clustered form. Clustered integrins form the adhesions to link the outside environments and inside cell signaling. The integrin-based adhesions are highly regulated to meet the requirement for rapid assembly and disassembly that happen within minutes. In this paper, we reported that kindlin2 as the core component in the FA, through oligomerization, undergoes phase separation with the clustered β-integrin in solution and on 2D membrane bilayer. Structure analysis identified that kindlin2 forms the oligomer through the intermolecular interaction between the PH domain and F0 lobe. The condensates formed by kindlin and integrin could also enrich other FA components. Further investigation in the cells showed that the kindlin2-dependent phase formation is essential for the adhesion formation, cell spreading, as well as integrin activation. Our discovery of the phase separation mediated by the oligomerized kindlin2 unveils the delicate mechanism to accommodate the rapid assembly of hundreds of the FA components to form the well-organized adhesions in minutes (Fig.7).

**Fig. 7.**
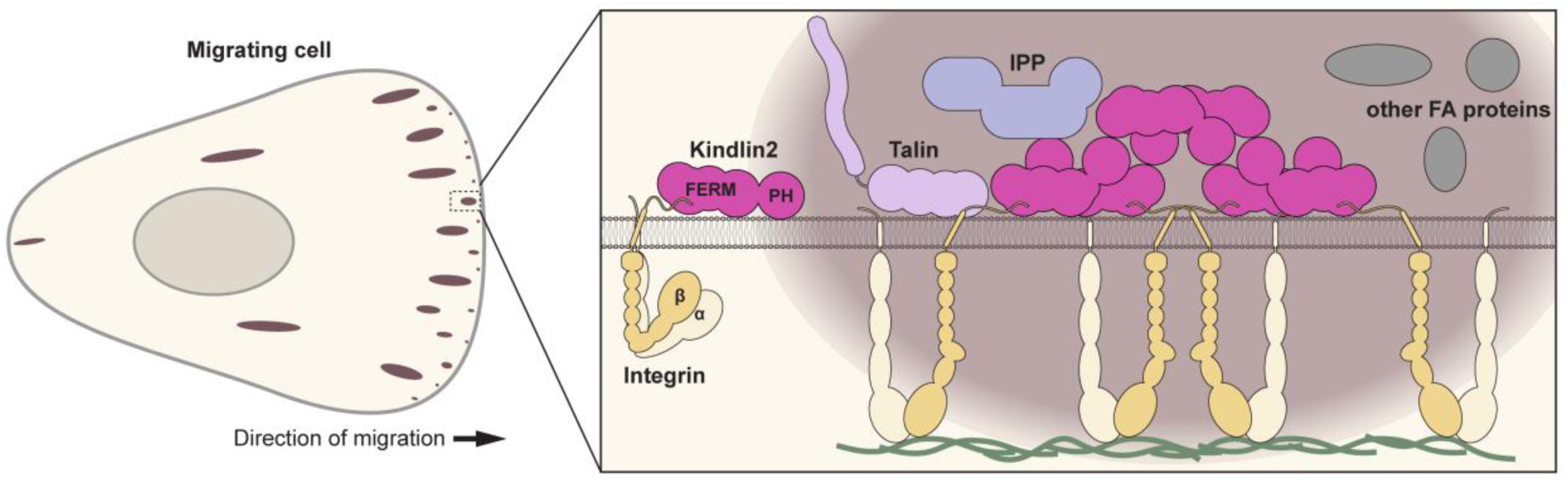
Proposed model of kindlin2 and integrin mediated phase separation in organizing focal adhesion formation. At the initial stage, kindlin2 binds to the integrin attached cell membrane as a monomer. In certain condition, kindlin2 undergoes conformational change between the FERM and PH domains to form dimer and to further form oligomer. The kindlin oligomer and clustered integrin form bio-condensates through phase separation, resulting in the enrichment of other FA components, including talin and the ILK/Parvin/PINCH (IPP) complex, to form the mature FA.

Recently, the understanding of the formation of membraneless organelles has revealed that many of these high dynamic assemblies form via liquid-liquid phase separation^34,39,49,50^. Multivalent weak interaction is an important way for generating phase separation in assembly of various membrane-associated compartments, such as immunological synapse^51,52^, presynaptic active zone^53^, post-synaptic density^35,37^, Tight-junctions^54,55^ and clustering of membrane receptors^36,56^. The advantage of the multivalent binding between the oligomerized kindlin2 and the clustered integrin is to drive the adhesion formation rapidly and dynamically. Consistently, the clustering of integrin has been reported to play important roles in adhesion formation^57,58^. Unlike the receptor clustering induced by phase separation in the previously reported membrane-associated compartments, our finding indicates that integrin clustering is the causation of the kindlin2-mediated phase formation, and the phase formation further promotes integrin clustering. Considering the widely existence of receptor clustering in various cellular processes, the phase separation mechanism by clustered integrins may be adopted by other clustered receptors.

The PH domain insertion in the FERM domain is unique for kindlin proteins. Despite that the importance of the PH domain for kindlins in integrin activation and ECM adhesion has been highly recognized by many studies^19,20,46-48,59^, the current understanding of the PH domain in kindlins is still limited and mainly relies on its phospholipid binding property, a common feature for PH domain-containing proteins. However, unlike other PH domains that only contains one C-terminal helix, the PH domain in kindlins contains a second C-terminal helix (α2^PH^, Extended Data Fig. 7a)^20^. Interestingly, the PH/F0 intermolecular interaction is mediated by the α2^PH^-helix (Fig. 4c). Mutating the highly conserved M498 in the α2^PH^-helix impairs the kindlin2-mediated phase separation in the 2D membrane system (Fig. 4f). Thus, other than the phospholipid binding, the PH domain employs the unique α2^PH^-helix to provide an intermolecular binding valency for the liquid-liquid phase separation in the integrin-based adhesion. As the inserted fragment in the F2 lobe, the PH domain is restricted in a close position to the F2 lobe, resulting in the intramolecular interaction between the PH domain and the F2 lobe (Extended Data Fig. 9a). Disrupting the PH/F2 interaction promotes the F2-based monomer-dimer transition of kindlin2 (Fig. 6d, Extended Data Fig. 8 and 10b), explaining at least in part the reason why the PH domain has to be inserted in the F2 lobe.

The PH/F0 intermolecular interaction is required for the phase separation on the lipid bilayer membrane but not for that in solution (Fig. 4e&f and Extended Data Fig. 3c&d). The weak PH/F0 interaction is likely to be stabilized in crystal due to the regular and linear packing between kindlin2 molecules (Fig. 4b), which is difficult to be achieved in solution. However, on the plasma membrane, once kindlins are attached, the flat 2D membrane facilitate the alignment of the PH domain to the favorable orientation for intermolecular packing. Consistently, in the cells transfected with kindlin2 containing the intermolecular interaction deficient mutant, the formed adhesions are smaller and easier to disappear than those cells transfected with the wild-type kindlin2 (Fig. 5f and Extended Data Fig. 6c, Movie S5). Recently, the F0 domain lobe and the PH domain has been reported to involved in the paxillin interaction^29,60,61^. Considering the importance of the PH/F0 interaction in the phase separation of kindlin, how the kindlin2 condensates are affected by paxillin will be further investigated.

The dimerization of kindlin2 is prerequisite for the phase separation. However, the *in vivo* factor(s) accelerating the monomer-dimer transition remain unclear. As an important regulator at the FA, ILK binds to kindlin2 through the loop connecting the N-terminus of the PH domain to the F2 lobe ^42^. As the intramolecular interaction between the loop in F2 domain and the PH domain inhibits the F2-based dimerization of kindlin2 (Fig. 6c and Extended Data Fig. 9), the ILK binding may disrupt this intramolecular interaction and therefore promote the kindin2 dimerization^62,63^, despite that we did not observe such a promotion effect *in vitro*. In addition, an IP_6_ molecule was found to interact with the PIP_3_-binding pocket of the PH domain in the kindlin2 dimer structure (Extended Data Fig. 7b). The addition of IP_6_ in the crystallization condition improved the crystal quality. Since the PIP_3_-binding loop in the PH domain is involved in the interdomain interaction between the PH domain and the dimerized F2 lobe (Extended Data Fig. 7b), the binding of IP_6_ to kindlin2 may enhance the dimer formation by stabilizing the PIP_3_-binding loop for the PH/F2 interaction. Interestingly, IP_6_ or its derivatives has been reported to regulate the FA-related cellular processes^64-66^.

## Methods and Materials

### Protein expression and purification

To obtain the kindlin2 full-length protein, mouse kindlin2 with a N-terminal His_6_-SUMO-tag was expressed in Rosetta (DE3) *E. coli* cells. The fusion protein was purified by Ni^2+^-NTA affinity chromatography followed by size-exclusion chromatography. The protein samples were then cleaved using SUMO protease and further purified by size-exclusion chromatography to remove tags. The di-kindlin2 protein was prepared as described previously^32^. In brief, the N-terminal truncated (residues 1 – 14) kindlin2 containing either the A318C or V522C mutation was purified. After removed SUMO tags, the two mutants were mixed at 1:1 ratio and incubated at 4 °C for 3 days. The mixture was then subjected to the analytical size-exclusion chromatography and the dimer fraction was collected for the experiments.

The mouse β1-integrin cytoplasm tail (β1-tail, residues 756 – 798) fused to a thioredoxin (Trx)-His_6_-GCN4 coiled-coil (250 – 281) tag was expressed in Rosetta (DE3) *E. coli* cells and purified by Ni^2+^-NTA affinity chromatography followed by size-exclusion chromatography. The tetrameric, trimeric, and hexameric GCN4 tag were designed based on the published papers^33^.

All point and truncation mutations of kindlin2 or β1-tail described in this study were prepared using PCR-based methods. The amino acid residues are labeled based on the kindlin2 isoform with 680 aa although the full length kindlin2 is the isoform containing 691 aa with additional 11 aa after G176 in the isoform 680. The mutant proteins were purified using essentially the same methods as those described for the wild-type protein.

ILK_KD (ILK kinase domain, residues 183 – 452)/Parvinα_CH2 (residues 248 – 372) complex was co-expressed in pET-Deut vector with the His_6_-tag fused to the N-terminus of ILK^43^ and was purified by Ni^2+^-NTA affinity chromatography followed by size-exclusion chromatography. Trx-talin_head (residues 1 – 400), Trx-kindlin2_PH (residues 367 – 512) and Trx-kindlin2_F2 (residues 260 – 572) with or without PH domain were purified essentially the same procedure as β1-tail. GST-kindlin2_F0 (residues 1 – 105) was purified by GST affinity chromatography followed by size-exclusion chromatography.

### Analytical gel filtration chromatography and multi-angle static light scattering

Analytical gel filtration chromatography was carried on an AKTA pure system (GE Healthcare). Protein samples were loaded onto a Superdex 200 Increase 10/300 GL column (GE Healthcare) equilibrated with the Protein Buffer (50 mM Tris, 100 mM NaCl, pH 7.5). Multi-angle static light scattering (MALS) assay was performed by coupling the analytical gel filtration chromatography system to a static light-scattering detector and differential refractive index detector (Wyatt). Data were analyzed with ASTRA6 provided by Wyatt.

### Isothermal titration calorimetry (ITC)

ITC measurements were carried out on a PEAQ-ITC Microcal calorimeter (Malvern) at 25 °C. All protein samples were dissolved in the Protein buffer. The titration processes were performed by injecting 3 μL aliquots of protein in the syringe into the protein in the cell. A time interval of 150 seconds between two titration points was applied to ensure that the titration peak returned to the baseline. The titration data were analyzed using PEAQ-ITC Analysis Software and plotted by OriginPro2020.

### Protein fluorescence labeling

For labeling, the purified proteins were pooled in PBS buffer (NaCl 137 mM, KCl 2.7 mM, Na_2_HPO_4_ 4.3 mM, KH_2_PO_4_ 1.4 mM, PH 8.3) and concentrated to 5∼10 mg/ml. Cy3/405 NHS ester (AAT Bioquest), Alexa Fluor 488 NHS aster (ThermoFisher), and Alexa Fluor 647 NHS aster (Invitrogen) were dissolved by DMSO and incubated with the corresponding protein at room temperature for 1 hour (molar ratio between fluorophore and protein was 1:1). Reaction was quenched by 200 mM Tris buffer, pH 7.5. The fluorophores and other small molecules were removed from the proteins by passing the reaction mixture through a Superdex 200 Increase 10/300 GL column (GE Healthcare) equilibrated with the Protein Buffer. In imaging assays, fluorescently labeled proteins were added into the corresponding unlabeled proteins in the same buffer with the final molar ratio of 1:50 (all the samples except for ILK) or 1:10 (ILK).

### *In Vitro* Phase Transition Assay

Kindlin2 and β1-integrin tail (wild-type or mutants) were prepared in the Protein Buffer. Typically, the two proteins were mixed at a 1:1 stoichiometry. For imaging, droplets were observed either by injecting mixtures into a homemade flow chamber or in the wells of the 96-well glass bottom plates. The homemade chamber was comprised of a glass slide covered by a coverslip with one layer of double-sided tape as a spacer for DIC (Differential Interference Contrast) and fluorescent imaging (Zeiss M2). The confocal images were taken from the samples in the 96-well glass bottom plates by Nikon A1R.

### *In vitro* FRAP Assay

20 μM di-kindlin and 20 μM tetra-β1CT were mixed in Protein Buffer containing additional 3 mM DTT. Sample was then diluted 4 times with the same buffer and loaded to a 96-well plate. FRAP assay was performed on a Nikon A1R Confocal Microscope at room temperature. Data was analyzed with Nikon NIS-Elements software.

### Lipid bilayer preparation

Phospholipids containing 95.9% POPC (Avantilipids), 4% DGS-NTA-Ni (Avantilipids) and 0.1% PEG 5000 PE (Avantilipids) were dried under a stream of nitrogen and resuspended by PBS to a final concentration of 0.5 mg/ml. The lipid solution was repeatedly frozen and thawed using a combination of liquid N2 and sonication until the solution turned clear. Then the solution was subjected to a centrifugation at 33,000 g for 45 min at 4 °C. Supernatant containing small unilamellar vesicles (SUVs) was collected.

The wells in 96-well plate (Lab-tek) was washed with 5M NaOH for 1 hour at 50 °C and thoroughly rinsed with ddH2O, repeated for three times, and followed by equilibration with the PBS Buffer. Typically, 50 μL SUVs were added to a cleaned well and incubated for 1 hour at 42 °C, allowing SUVs to fully collapse on glass and fuse to form supported lipid bilayers (SLBs). SLBs were washed with the Protein Buffer for three times (5-folds dilution per time, 125-folds dilution in total) to remove extra SUVs. The SLB was blocked with the Cluster Buffer (the Protein Buffer supplied with 1 mg/ml BSA) for 30 minutes at room temperature.

### Lipid bilayer-based phase transition assay

The supported membrane bilayers were doped with DGS lipid with Ni^2+^-NTA attached to its head. The tetra-β1CT fused with an N-terminal Trx-His_6_ tag, and the fluorescence tag (Alexa Fluor 488 in this study) can be added to the N-terminus of the fusion protein (on the Trx N-terminus) without affecting the His_6_-tag from attaching to Ni^2+^-NTA-DGS embedded in the lipid bilayer. The Trx-tag also provides a space to separate the chemical fluorophore from the surface of lipid bilayers to avoid potential non-specific interaction between the dye and the membranes. Initially, 50 μM His-tetra-β1CT was added and incubated with SLBs for 1 hour at room temperature, followed by washing with the Cluster Buffer for three times (5-folds dilution per time) to remove unbound His-tetra-β1CT. Kindlin or other proteins were added to the His-tetra-β1CT-bound SLBs, waiting for phase transition to happen on the lipid bilayers. All data were collected within 8 hours after lipid coating started.

### Crystallization

To determine the crystal structure of the full-length kindlin, various constructs of mouse kindlin2 containing the deletion of the flexible loops in the N-terminus and the F1 lobe was expressed and purified. Crystals were obtained by the sitting drop vapor diffusion method at 16 °C. To set up a sitting drop, 1 μL 30 mg/ml protein solution was mixed with 1 μL of crystallization solution with 2% v/v Tacsimate pH 7.0, 5% v/v 2-Propanol, 0.1 M Imidazole pH 7.0, 8% w/v Polyethylene glycol 3,350 for the kindlin2 protein with the deletion of residues 1 – 14 and 168 – 217 (kindlin2-FL1). To optimize the crystal, 0.2 μL additive with 0.16% w/v L-Lactic acid, 0.16 % w/v L-Aspartic acid, 0.16% w/v L-Thyroxine, 0.16% w/v Pyridoxine, 0.16% w/v L-Ascorbic acid, 0.16% w/v Phytic acid (IP_6_) sodium salt hydrate, 0.02M HEPES sodium pH 6.8 was added in the drops. The crystal of kindlin2 with the deletion of residues 1 – 14 and 149 – 204 (kindlin2-FL2) were grown in a precipitant containing 0.5 M Bis-Tris propane pH 7.0, 2M Sodium acetate trihydrate.

### Diffraction data collection and structure determination

Before X-ray diffraction experiments, crystals were soaked in crystallization solution containing 30% glycerol for cryoprotection. The diffraction data were collected at Shanghai Synchrotron Radiation Facility beamline BL17U^67^, BL18U and BL19U1. The data were processed and scaled using the HKL2000^68^. The dimeric and monomeric kindlin2ΔPH structures (PDB id: 5XPZ and 5XPY) were used as the search models for phase determination of the two kindlin2 crystals by molecular replacement. Kindlin2-FL1 and kindlin2-FL2 were solved as the dimer and monomer structures, respectively. IP_6_ molecules, existing in the crystallization additive, were further built into the model of kindlin2-FL1. These models were refined again in PHENIX^69^. COOT was used for model rebuilding and adjustments^70^. In the final stage, an additional TLS refinement was performed in PHENIX. The model quality was check by MolProbity ^71^. The final refinement statistics are listed in Table 1. All structure figures were prepared by PyMOL (http://www.pymol.org/).

### Cell culture and transfection

Human HT1080 kindlin2 knockout cells were cultured at 37 °C with 5% CO_2_ in MEM supplemented with 10% FBS, 0.1mM non-essential amino acids and 50 U/ml penicillin and streptomycin. CHO-A5 cells stably expressing integrin αIIbβ3 were cultured at 37 °C with 5% CO_2_ in DMEM/F12 supplemented with 10% FBS, 0.1mM non-essential amino acids, 50 U/ml penicillin and streptomycin. Transfection was performed using Lipofectamine 3000 according to manufacturer’s instructions.

### Confocal microscopy

24 hours after transfection, cells were resuspended and re-plated on coverslips coated with 20 μg/ml fibronectin. 2 hours after re-plating, cells were washed once with PBS and then fixed at 37 °C for 10 minutes in 4% paraformaldehyde. Fixed cells were incubated in blocking buffer (2% BSA and 0.1% Triton X-100 in PBS) at room temperature for 15 minutes, then stained with anti-paxillin antibody (1:200) in the same buffer at 4 °C overnight followed by incubation with Alexa Fluor 594 anti-mouse IgG Ab (1:1000, Invitrogen) for 1 hour at room temperature. Images were taken using Nikon A1R confocal microscope with 100X oil immersion objective. Images were analyzed using ImageJ software.

### Total internal reflection fluorescence microscope (TIRFM)

Cells were trypsinized and resuspended 24 hours after transfection, and then added to glass bottom cell culture dish coated with 20 μg/ml fibronectin for live TIRF imaging. All TIRF images were taken within 1 hour after cells were added to the dishes.

### Cell spreading assay

24 hours after transfection, cells were sorted by GFP fluorescence using FACS sorter (BD FACS AriaIII). Sorted cells were then seeded on fibronectin coated coverslip and incubated at 37 °C for 2 hours. Cells were then fixed and stained with Alexa Fluor 594 Phalloidin for F-actin and observed with Leica DMI6000B fluorescence microscope using a 10X objective. Image was analyzed by ImageJ software.

### Integrin activation assay

Integrin activation assay was done as previously described^32^. Briefly, CHO-A5 cells stably expressing αIIbβ3-integrin were transfected with GFP-tagged kindlin2 (wild-type or mutants) and RFP-tagged talin-FERM. 24 hours after the transfection, cells were collected and incubated with mAb PAC-1 (10μg/ml) for 30 minutes followed by staining with Alexa Fluor 633-conjugated goat anti-mouse IgM Ab (1:1000, Invitrogen) for 30 minutes at room temperature. Cells were then fixed with 4% polyformaldehyde for 10 minutes and analyzed using a FACSCantoTM (BD) flow cytometer. PAC1-binding was analyzed on a gated subset of double-positive cells. Integrin activation was expressed in terms of relative median fluorescence intensities by normalizing the basal PAC1-binding in control cells (CHO-A5) to 1.0. Three or four independent experiments were performed.

## Acknowledgements

We thank Drs. Wenjie Wei, Yilin Wang and Pengfei Li for their suggestions on the experiment design and Dr. Xingqiao Xie and Dr. Yuqun Xu for their technical support on the sample preparation. We thank the assistance of Southern University of Science and Technology (SUSTech) Core Research Facilities. We thank the staff from BL17U, BL18U and BL19U1 beamlines of Shanghai Synchrotron Radiation Facility for assistance during data collection. This work was supported by the National Natural Science Foundation of China (Grant No. 31870757 to C.Y., 31971131 and 31770791 to Z.W.), Science and Technology Planning Project of Guangdong Province (2017B030301018), Shenzhen-Hong Kong Institute of Brain Science, Shenzhen Fundamental Research Institutions (2019SHIBS0002), and Funds for High Level Universities (G02226301). Z.W. is a member of the Brain Research Center, SUSTech.

## Author contributions

C.Y. conceived the study. C.Y. and Z.W. co-supervised the project. Y.L., H.Y., and H.L. designed constructs. Y.L., H.Y., R.L., L.W. and J.Z. purified proteins. Y.L. and T.Z. performed biochemistry assays. H.L. and Z.W. solved and analyzed structures. T.Z., C.Y., and K.S. designed cellular experiments. T.Z. performed cellular assays. T.Z., K.S., C.Y., and Z.W. analyzed the cellular data. C.Y. and Z.W. wrote the manuscript.

## Figures

**Extended Data Fig. 1.**
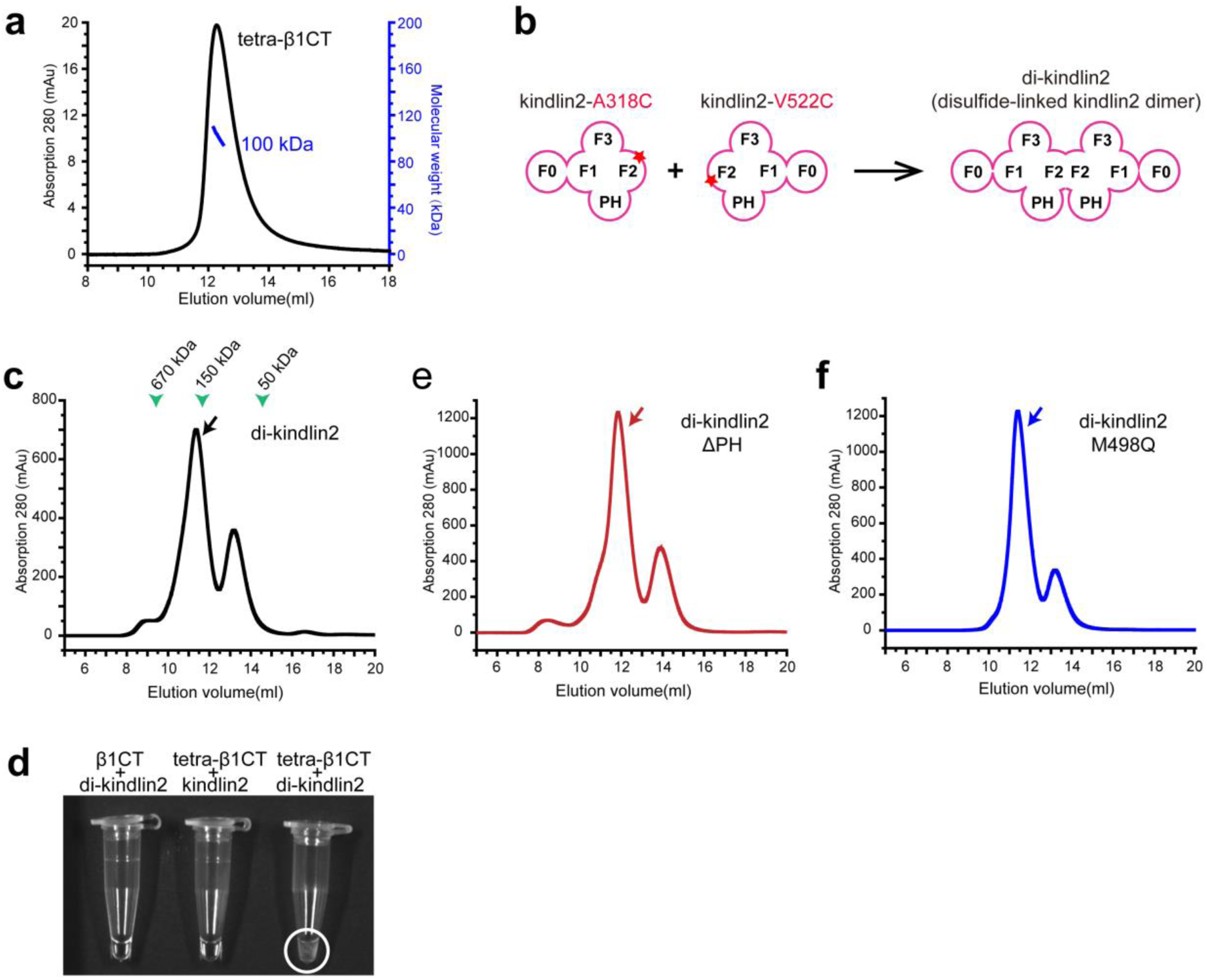
Purification of clustered-β1CT and dimerized-kindlin. **a**, The molecular weight of purified GCN4-tetramerized β1-integrin was 100 kDa, as measured by Multi-angle static light scattering. The theoretical molecular weight of monomer is 24 kDa. **b**, Schematics of di-kindlin2 formation. **c**, Gel filtration analysis of di-kindlin2 WT. The eluted kindlin2 dimer peaks of wild-type and mutants are indicated by an arrow in each panel. **d**, Mixing tetra-β1CT and di-kindlin2 resulted in an opalescent solution (as indicated by the white circle) while β1CT monomer and di-kindlin2 or tetra-β1CT and kindlin2 monomer remained clear. **e&f**, Gel filtration analysis of di-kindlin2 variants, ΔPH (**e**) and M498Q (**f**).

**Extended Data Fig. 2.**
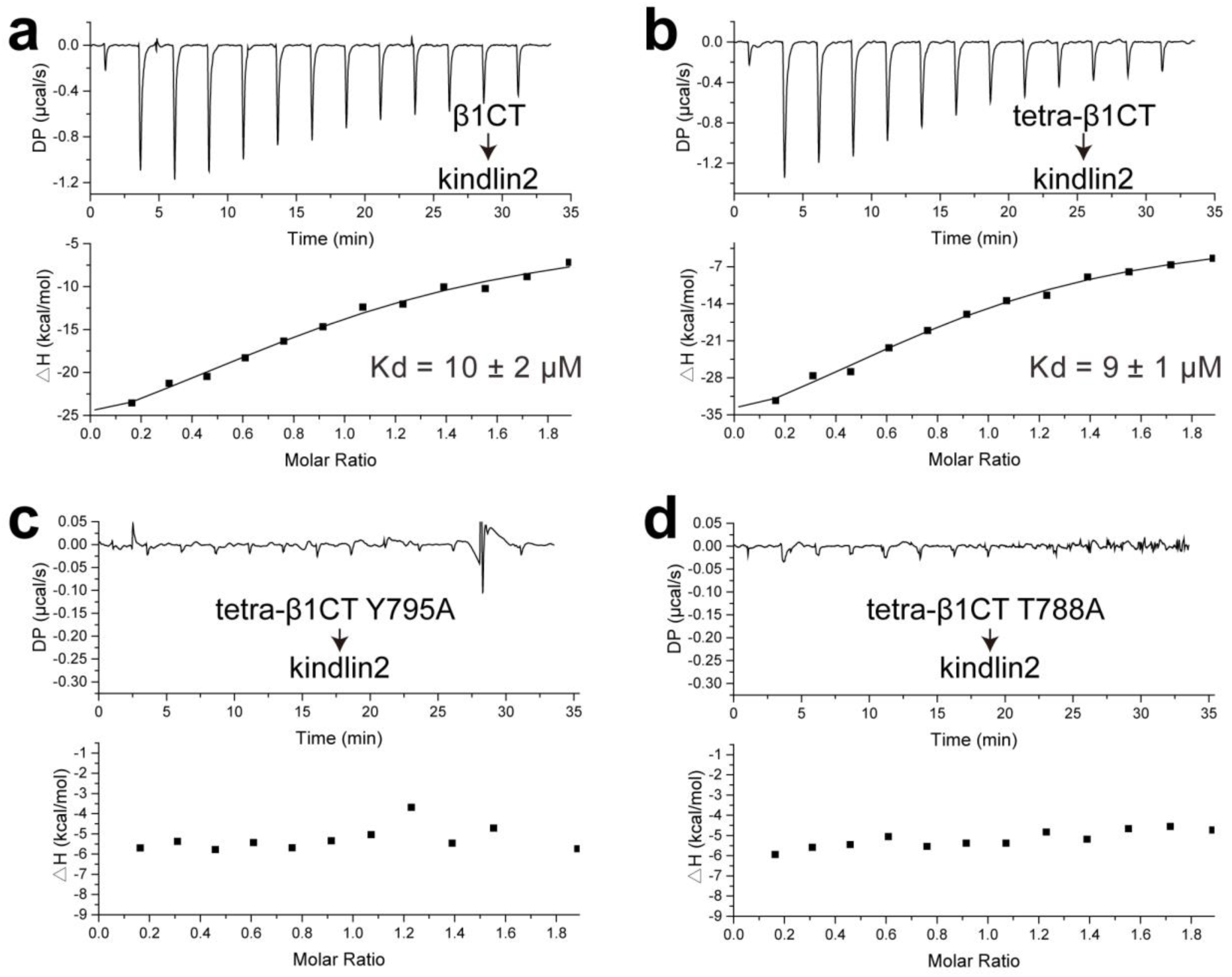
ITC analysis of the interactions between kindlin2 and β1-integrin tail. ITC curves show the various forms of β1-tail (200 μM) titrated to kindlin2 (20 μM). *K*_d_ value with standard error of mono-(**a**) or tetra-(**b**) β1CT was indicated. The Y795A (**c**) and T788A (**d**) mutants of tetra-β1CT show no detectable binding to kindlin2.

**Extended Data Fig. 3.**
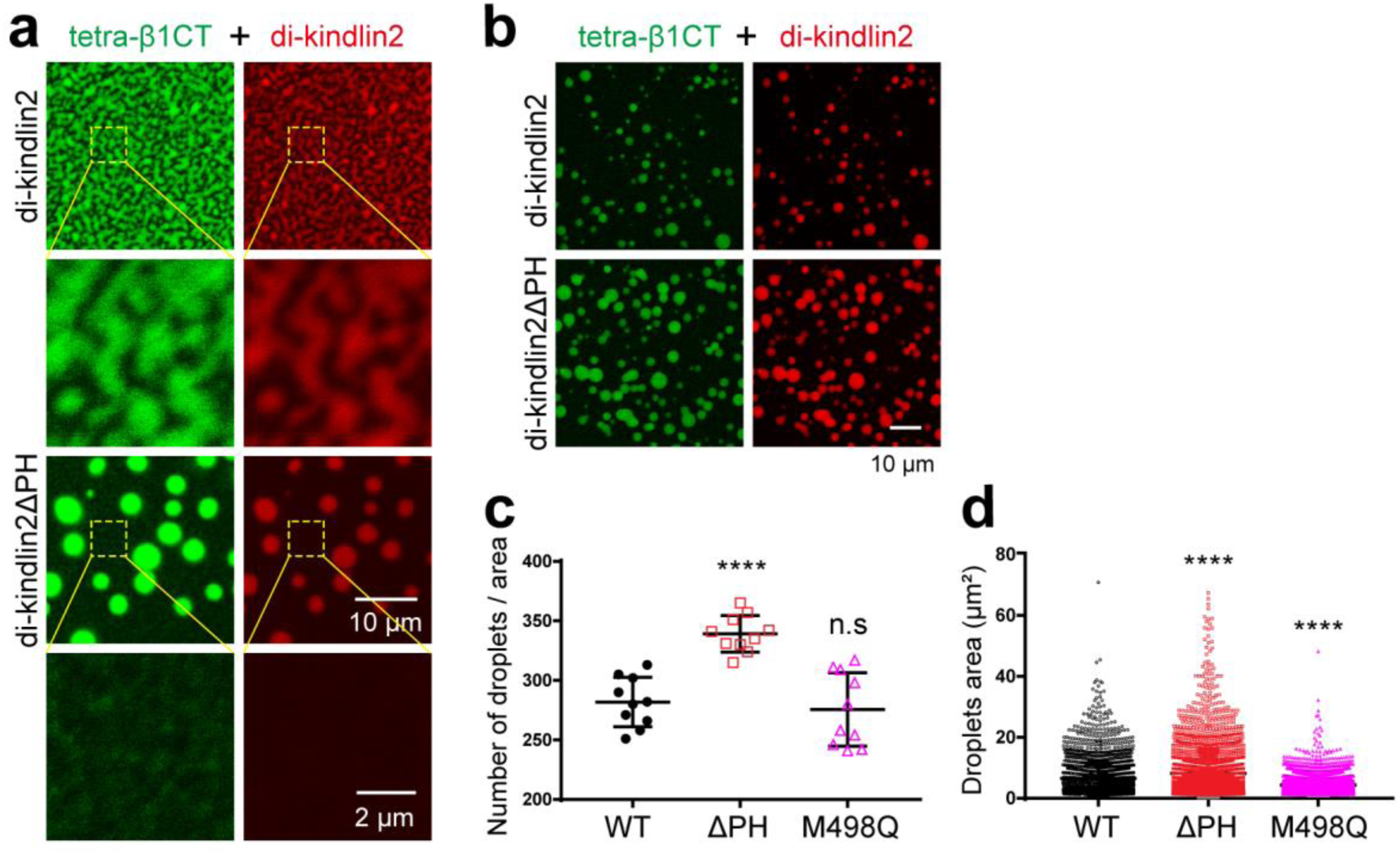
Deletion of the PH domain in kindlin2 diminishes the kindlin2/integrin condensates formation on the membrane but not in solution. **a**, 2 μM of di-kindlin2 or di-kindlin2 with the PH domain deletion mutation (di-kindlin2ΔPH) was added to the SLB with His-tetra-β1CT, shown by confocal microscopy. **b**, Di-kindlin2 or di-kindlin2ΔPH and tetra-β1CT form condensates in solution, shown by confocal microscopy. The concentration of each protein was 20 μM, and the mixture was diluted 4 times with the same buffer for observation. **c**, Quantification of the condensates numbers as shown in the panel b and Fig. 4e. Confocal microscopy images were taken 20 minute later and 10 images (image area 15099 μm^2^) from each sample were analyzed for quantification. **d**, Quantification of the condensates area as shown in panel b and Fig. 4e. In each condition, 10 images of different views were analyzed by measuring the size and number of droplets using ImageJ.

**Extended Data Fig. 4.**
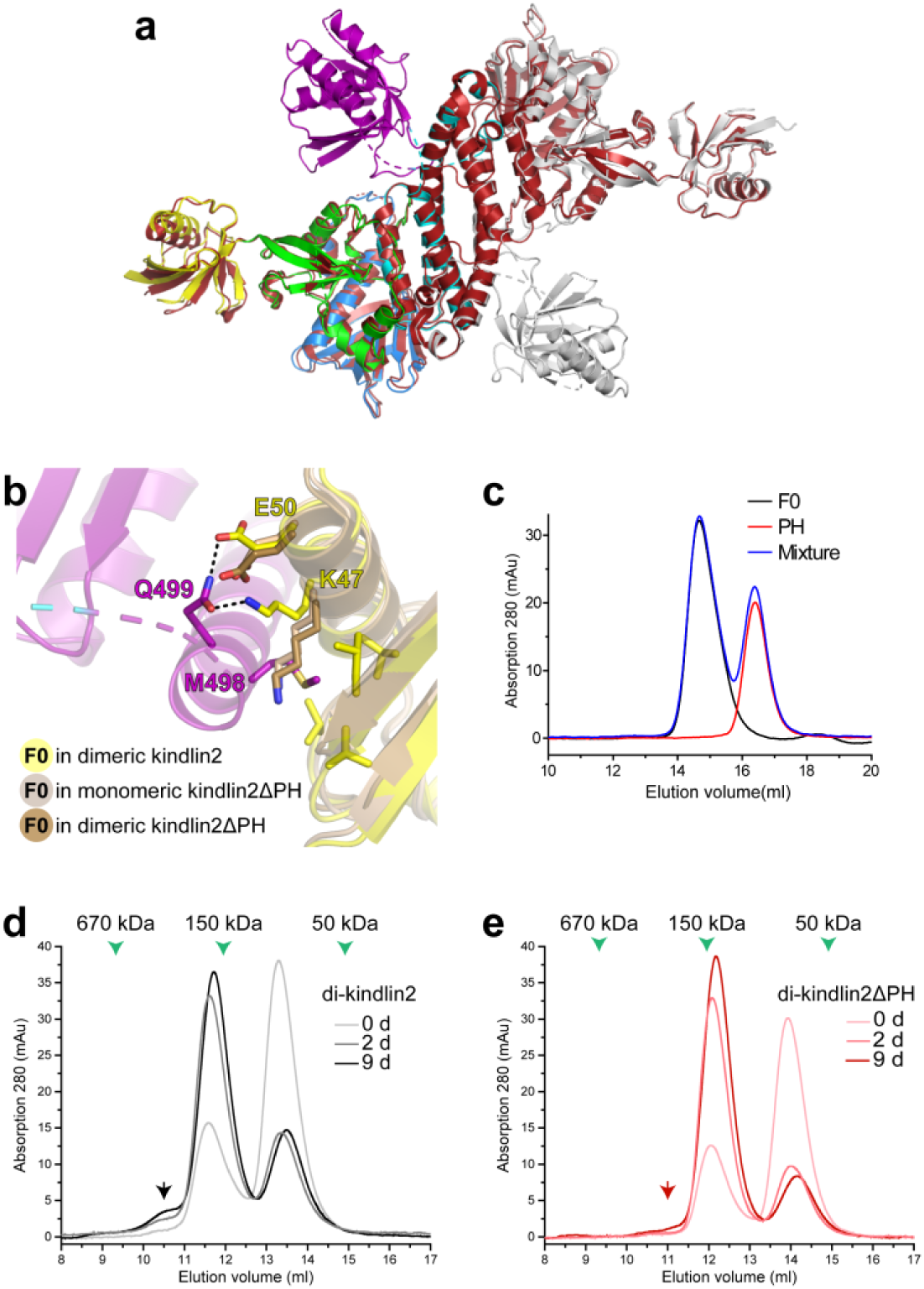
Structural and biochemical analysis of the PH/F0 interaction. **a**, Structure comparison of the kindlin2 and kindlin2ΔPH dimers. The kindlin2ΔPH (PDB id: 5XPZ) dimer was colored in red. **b**, Sidechain conformational changes of K47 and E50 in the F0 lobe. **c**, Analytical gel filtration analysis of the weak F0/PH interaction. No complex formation was observed by using the mixture of the GST-tagged F0 lobe and the Trx-tagged PH domain at concentration of 20 μM. **d**, Detection of the kindlin2 oligomerization in solution. The A318C and V522C mutants of kindlin2 each at the concentration of 300 μM were mixed to prepare the sample stock. At different time point (0, 2, and 9 days after mixing), the stock was aliquot and diluted to 20 μM and subjected to analytical gel filtration. The oligomer fraction was indicated by a dark arrow. **e**, Removal of the PH domain from kindlin2 blocking the oligmerization. The same strategy described in the panel **d** was applied to analyze the di-kinldin2ΔPH sample. The corresponding oligomer fraction was indicated by a red arrow.

**Extended Data Fig. 5.**
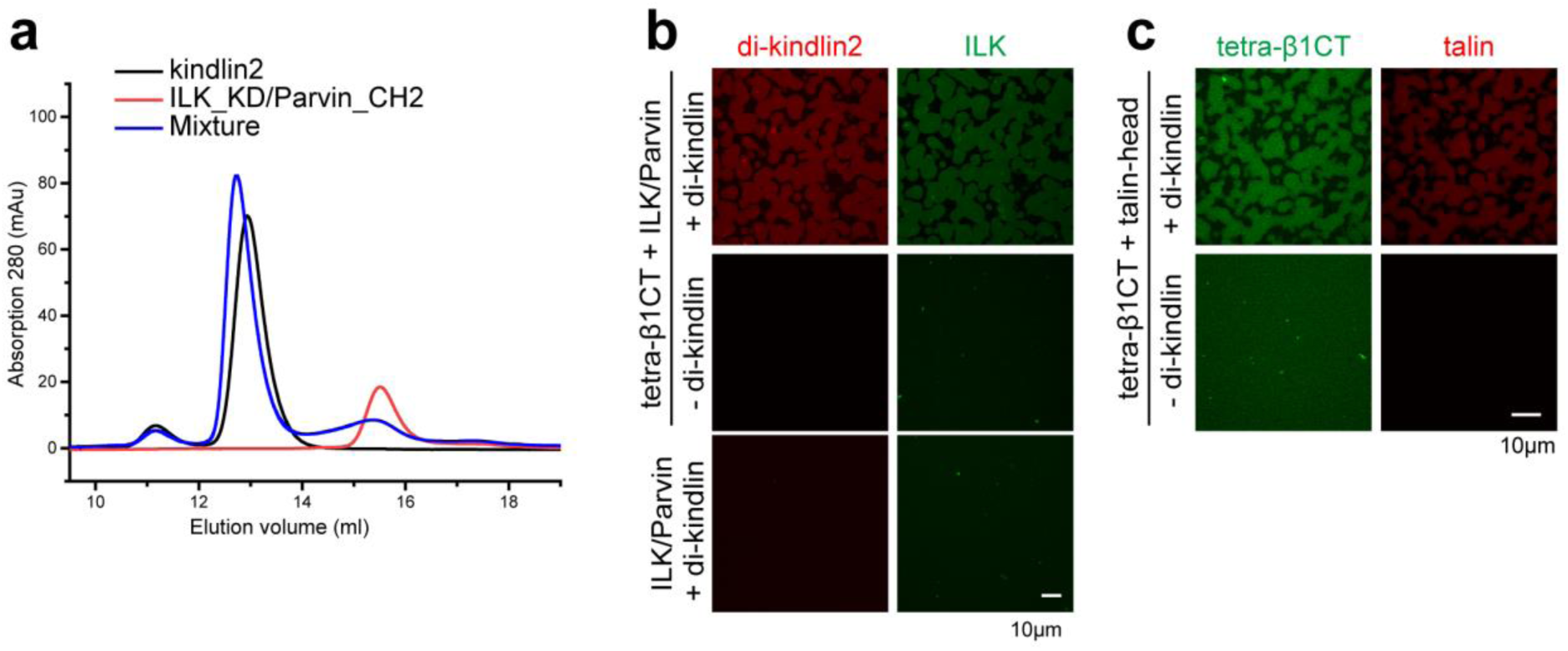
FA components enter the condensates formed by kindlin2 and integrin in solution. **a**, Analytical gel filtration analysis of the interaction between kindlin2 and the ILK_KD/Parvin_CH2 complex. The sample concentration was 20 μM for SUMO-kindlin2 and the His-ILK_KD/Parvin_CH2 complex. **b**, Confocal imaging of liquid droplets formed by mixing di-kindlin2, tetra-β1CT and the ILK_KD/Parvin_CH2 complex at 20 μM each. Fluorescence analysis showed that the ILK complex failed to undergo phase separation with di-kindlin2 or tetra-β1CT (middle and bottom), whereas the complex entered the kindlin2/integrin condensates (Top). **c**, Similar with ILK, talin_head also entered the kindlin2/integrin condensates at the concentration of 40 μM.

**Extended Data Fig. 6.**
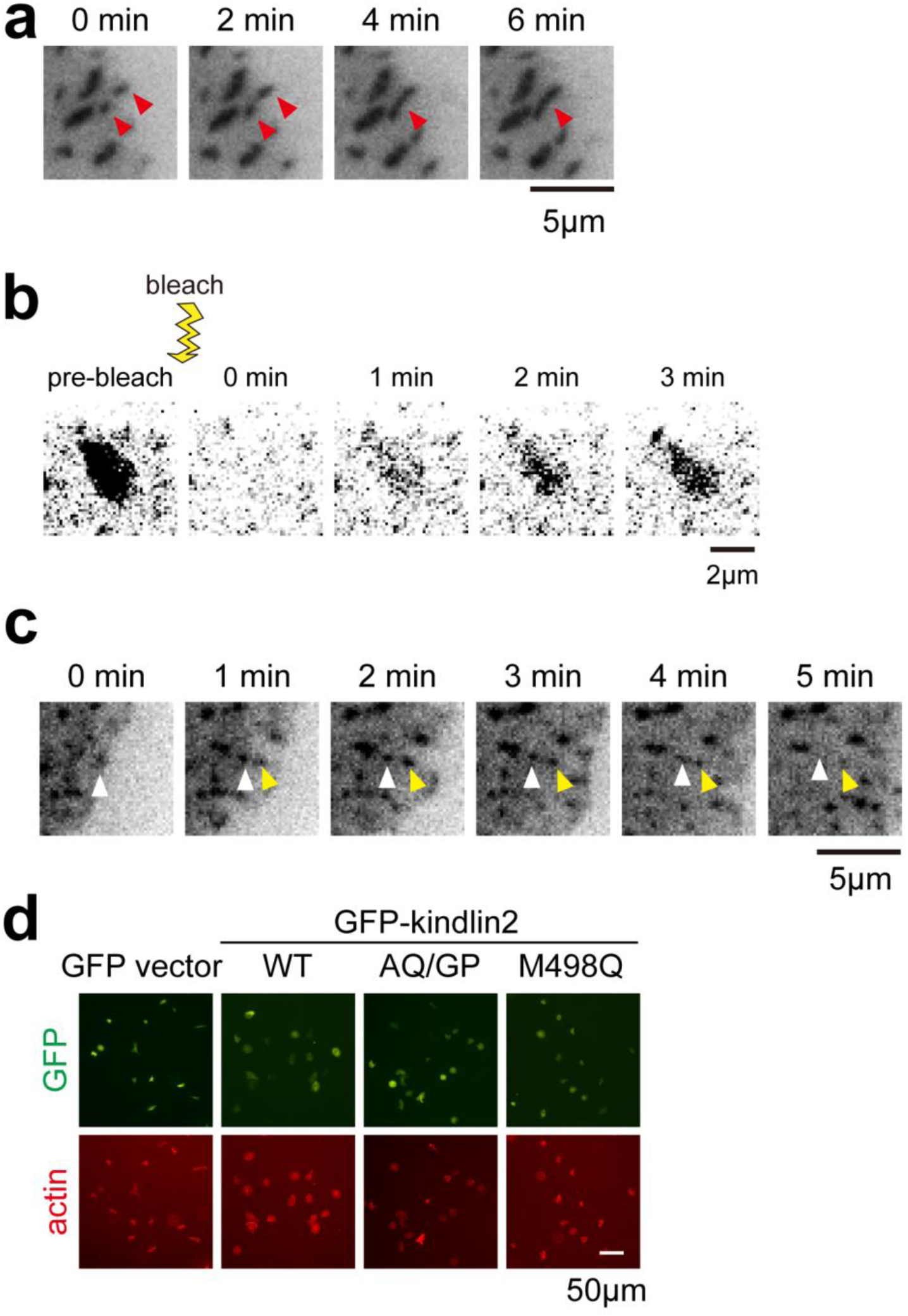
The dynamic property of focal adhesions in cells. **a**, The FAs (indicated by red arrowhead) fused with each other as indicated by the GFP-kindlin2 signaling taken by TIRF microscopy. **b**, FRAP analysis of GFP-kindlin2 at the FA by confocal microscopy. **c**, Rapid adhesions turnover in the kindlin2-knockout HT1080 cells transfected with GFP-kindlin2_M488Q showing by TIRF microscopy. Two adhesions, as indicated by arrowhead, disappeared quickly. **d**, Cell spreading analysis of the kindlin2 mutants. Kindlin2-KO cells were transfected with GFP vector or GFP-tagged kindlin2 variants. Cell areas were indicated by actin staining. The quantified results were indicated in Fig.**5e**.

**Extended Data Fig. 7.**
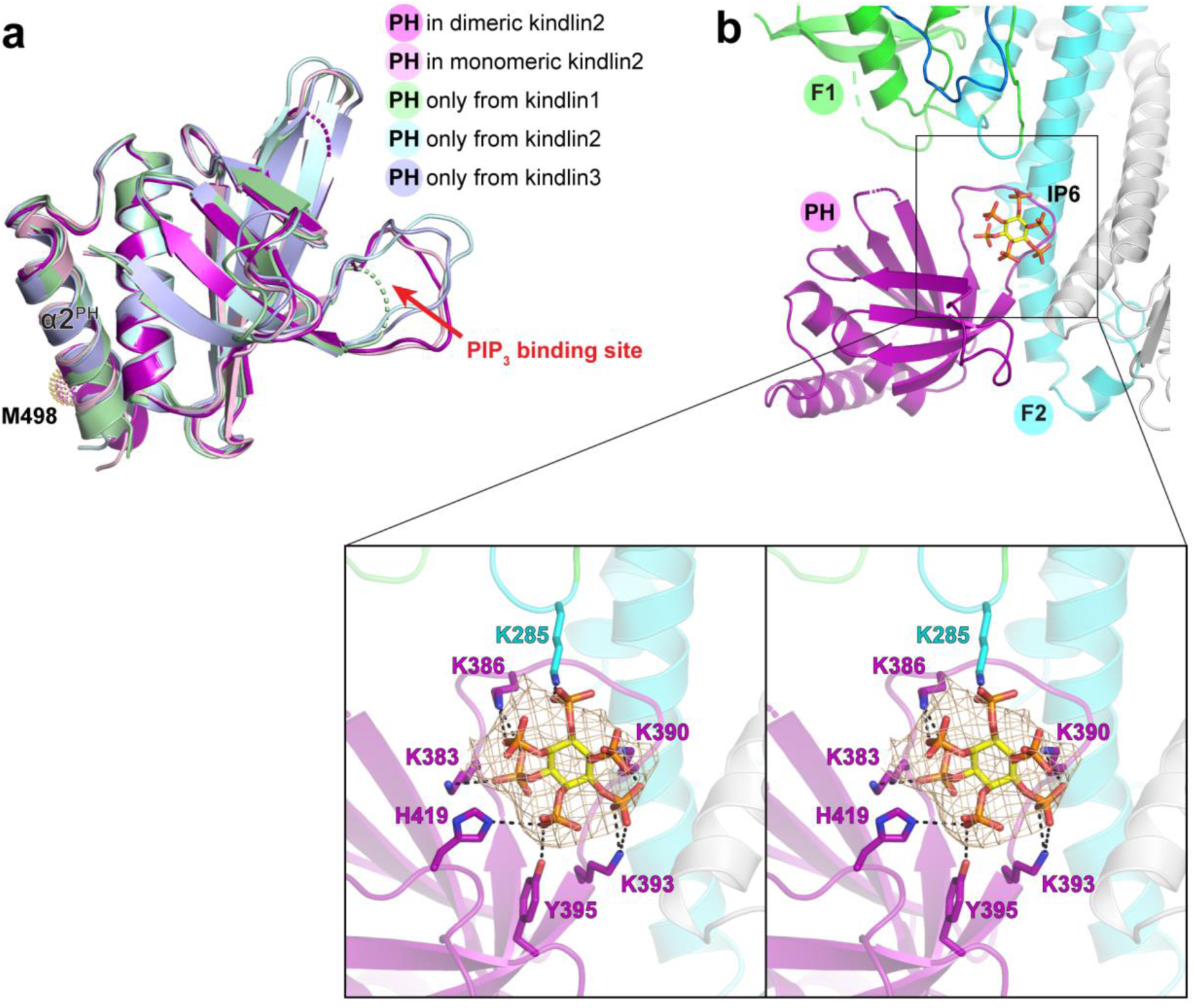
Structure analysis of the PH domains and their PIP_3_-binding pockets. **a**, Structural comparison of the solved PH domain structures in kindlins, including kindlin1_PH (PDB id: 4BBK), kindlin2_PH (4F7H), and kindlin3_PH (5L81). The PIP_3_-binding site was indicated by a red arrow. M498 locates on the α2^F1^-helix that is far away from the PIP_3_-binding site. **b**, An IP_6_ molecule was modeled into the electron density observed in the PIP_3_-binding pocket of the kindlin2 dimer. Hydrogen bonds and salt bridges were indicated by dash lines.

**Extended Data Fig. 8.**
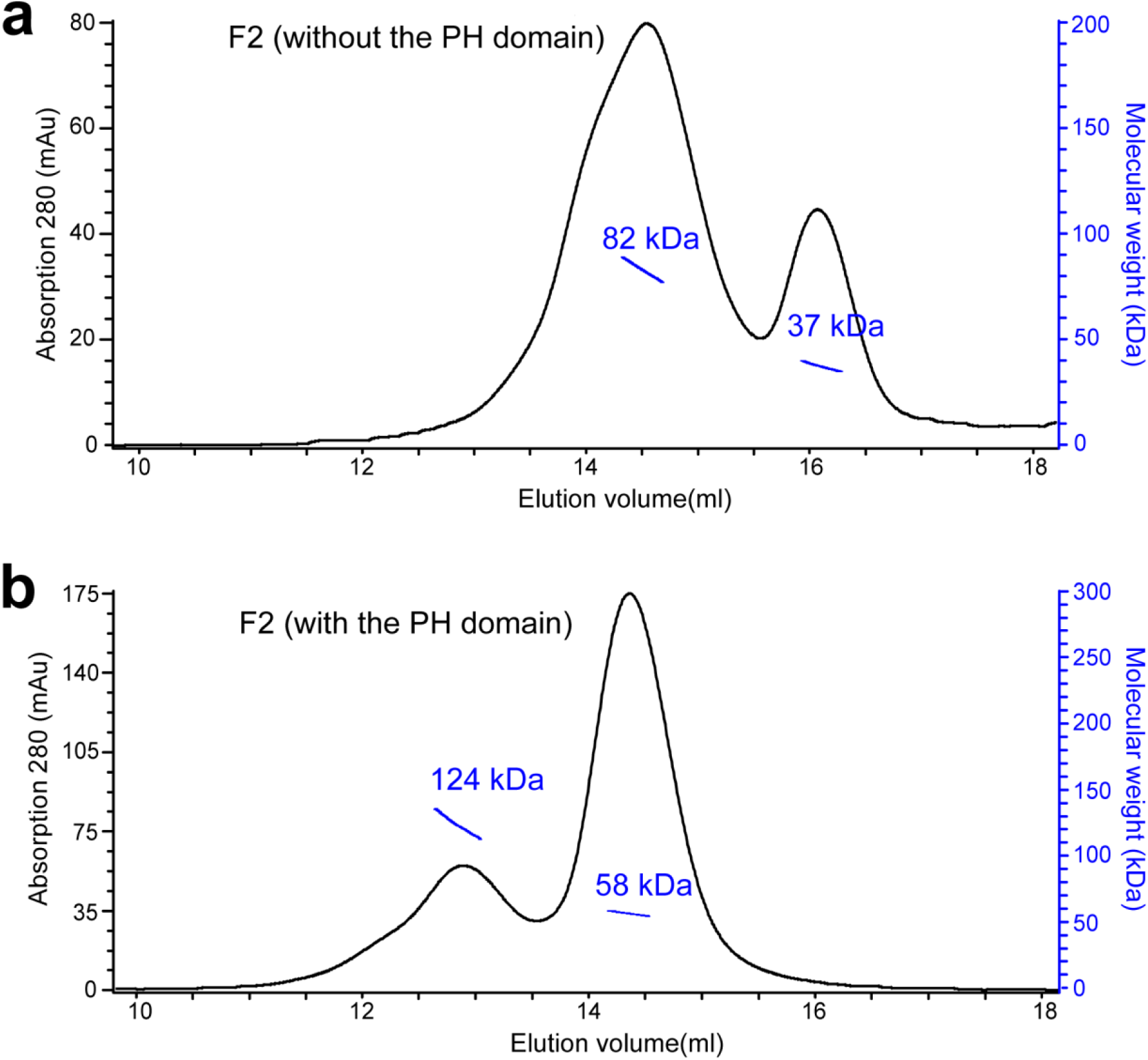
The F2 lobe forms a stable dimer. **a**, The purified Trx-F2 (without the PH domain) is predominantly dimer, as measured by multi-angle static light scattering. The theoretical molecular weight is 34 kDa. **b**, The dimer fraction is largely decreased in the purified Trx-F2 (with the PH domain). The theoretical molecular weight is 51 kDa.

**Extended Data Fig. 9.**
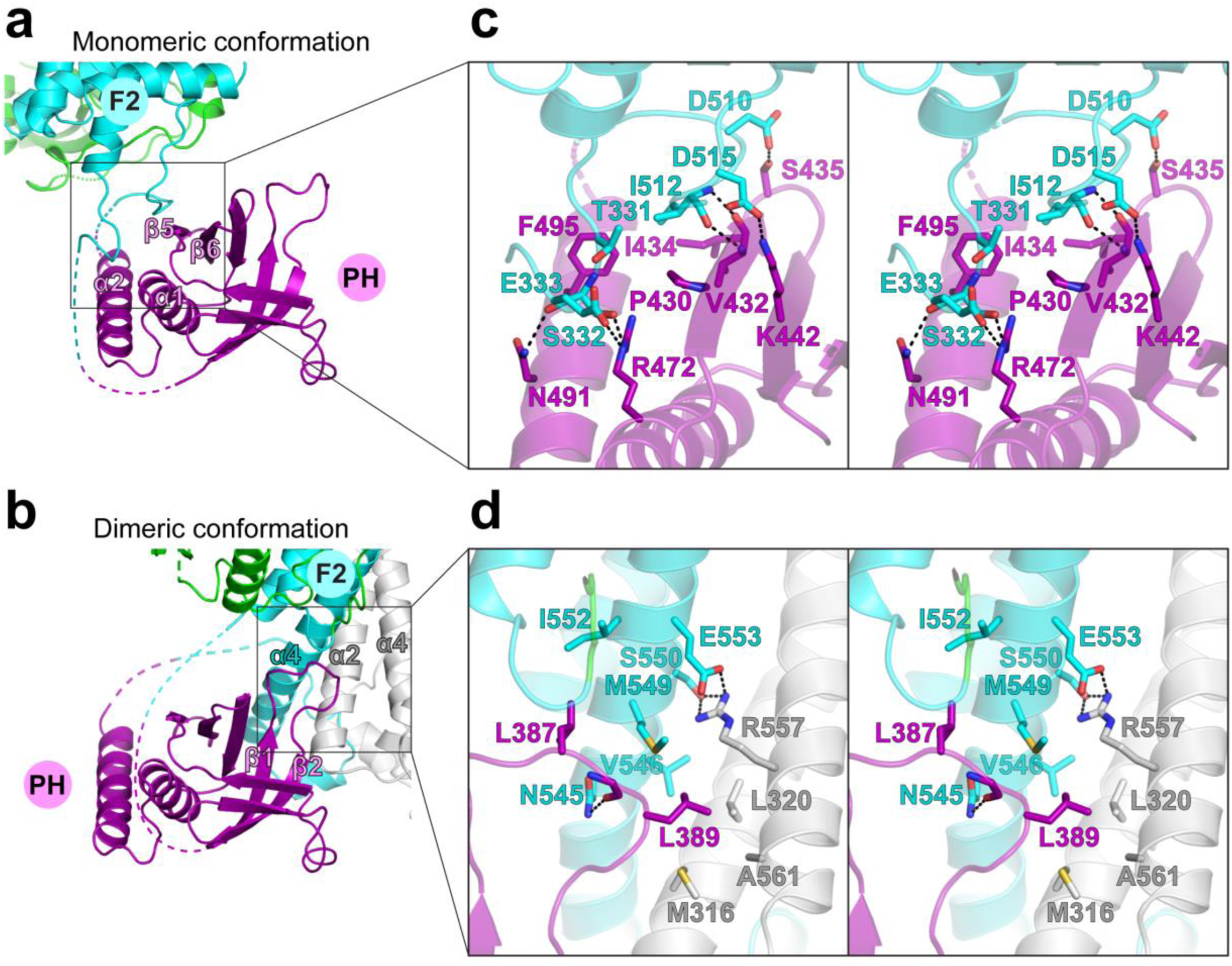
Structural analysis of the PH/F2 interfaces in the monomeric (a) and dimeric (b) conformations of kindlin2. The molecular details of the interfaces were shown as stereoviews in panel **c** and **d**. Hydrogen bonds and salt bridges were indicated by dotted lines. The PIP_3_-binding loop directly interacts with the dimer interface in the kindlin2 dimer.

**Extended Data Fig. 10.**
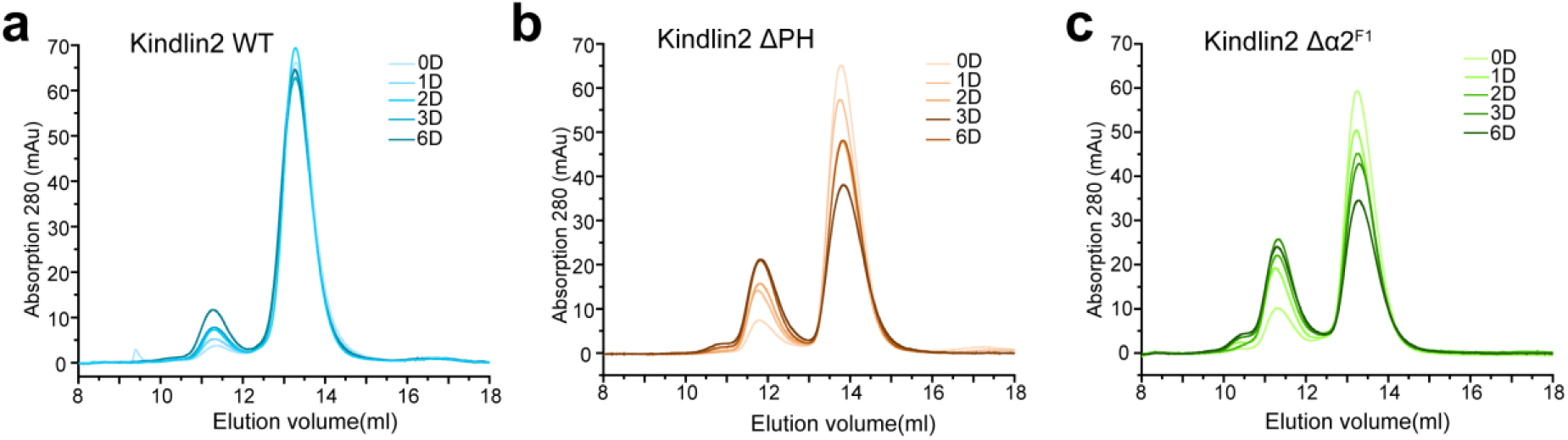
Time-dependent dimerization analysis of kindlin2 variants. The samples at the concentration of 30 μM were prepared. At different time point (0, 1, 2, 3, and 6 days), the samples were subjected to analytical gel filtration.

Movie S1. Related to Extended Data Fig. 6a, the FAs (indicated by red arrowhead) fused with each other as indicated by the GFP-kindlin2 signaling taken by TIRF microscopy.

Movie S2. Related to Fig. 5c, TIRF live cell imaging of adhesion formation in the kindlin2-KO cells transfected with GFP-kindlin2 wild-type.

Movie S3. Related to Fig. 5c, TIRF live cell imaging of adhesion formation in the kindlin2-KO cells transfected with GFP-kindlin2 AQ/GP.

Movie S4. Related to Fig. 5c, TIRF live cell imaging of adhesion formation in the kindlin2-KO cells transfected with GFP-kindlin2 M498Q.

Movie S5. Related to Extended Data Fig. 6a, rapid adhesions (indicated by red arrowhead) turnover in the kindlin2-knockout HT1080 cells transfected with GFP-kindlin2_M488Q showing by TIRF microscopy.

## References

1 Geiger, B., Bershadsky, A., Pankov, R. & Yamada, K. M. Transmembrane crosstalk between the extracellular matrix--cytoskeleton crosstalk. Nature reviews. Molecular cell biology 2, 793–805 (2001).

2 Ginsberg, M. H., Partridge, A. & Shattil, S. J. Integrin regulation. Current opinion in cell biology 17, 509–516 (2005).

3 Hynes, R. O. Integrins: bidirectional, allosteric signaling machines. Cell 110, 673–687 (2002).

4 Winograd-Katz, S. E., Fassler, R., Geiger, B. & Legate, K. R. The integrin adhesome: from genes and proteins to human disease. Nature reviews. Molecular cell biology 15, 273–288 (2014).

5 Seguin, L., Desgrosellier, J. S., Weis, S. M. & Cheresh, D. A. Integrins and cancer: regulators of cancer stemness, metastasis, and drug resistance. Trends in cell biology 25, 234–240 (2015).

6 Iwamoto, D. V. & Calderwood, D. A. Regulation of integrin-mediated adhesions. Current opinion in cell biology 36, 41–47 (2015).

7 Zaidel-Bar, R., Itzkovitz, S., Ma’ayan, A., Iyengar, R. & Geiger, B. Functional atlas of the integrin adhesome. Nature cell biology 9, 858–867 (2007).

8 Gardel, M. L., Schneider, I. C., Aratyn-Schaus, Y. & Waterman, C. M. Mechanical integration of actin and adhesion dynamics in cell migration. Annual review of cell and developmental biology 26, 315–333 (2010).

9 Bachir, A. I. et al. Integrin-associated complexes form hierarchically with variable stoichiometry in nascent adhesions. Current biology : CB 24, 1845–1853 (2014).

10 Harburger, D. S. & Calderwood, D. A. Integrin signalling at a glance. J Cell Sci 122, 159–163 (2009).

11 Morse, E. M., Brahme, N. N. & Calderwood, D. A. Integrin cytoplasmic tail interactions. Biochemistry 53, 810–820 (2014).

12 Horton, E. R. et al. Definition of a consensus integrin adhesome and its dynamics during adhesion complex assembly and disassembly. Nature cell biology 17, 1577–1587 (2015).

13 Tu, Y., Wu, S., Shi, X., Chen, K. & Wu, C. Migfilin and Mig-2 link focal adhesions to filamin and the actin cytoskeleton and function in cell shape modulation. Cell 113, 37–47 (2003).

14 Shi, X. et al. The MIG-2/integrin interaction strengthens cell-matrix adhesion and modulates cell motility. The Journal of biological chemistry 282, 20455–20466 (2007).

15 Ma, Y. Q., Qin, J., Wu, C. & Plow, E. F. Kindlin-2 (Mig-2): a co-activator of beta3 integrins. The Journal of cell biology 181, 439–446 (2008).

16 Montanez, E. et al. Kindlin-2 controls bidirectional signaling of integrins. Genes & development 22, 1325–1330 (2008).

17 Moser, M., Nieswandt, B., Ussar, S., Pozgajova, M. & Fassler, R. Kindlin-3 is essential for integrin activation and platelet aggregation. Nature medicine 14, 325–330 (2008).

18 Ye, F. et al. The mechanism of kindlin-mediated activation of integrin alphaIIbbeta3. Current biology : CB 23, 2288–2295 (2013).

19 Qu, H. et al. Kindlin-2 regulates podocyte adhesion and fibronectin matrix deposition through interactions with phosphoinositides and integrins. J Cell Sci 124, 879–891 (2011).

20 Liu, J. et al. Structural basis of phosphoinositide binding to kindlin-2 protein pleckstrin homology domain in regulating integrin activation. The Journal of biological chemistry 286, 43334–43342 (2011).

21 Meves, A., Stremmel, C., Gottschalk, K. & Fassler, R. The Kindlin protein family: new members to the club of focal adhesion proteins. Trends in cell biology 19, 504–513 (2009).

22 Larjava, H., Plow, E. F. & Wu, C. Kindlins: essential regulators of integrin signalling and cell-matrix adhesion. EMBO reports 9, 1203–1208 (2008).

23 Jobard, F. et al. Identification of mutations in a new gene encoding a FERM family protein with a pleckstrin homology domain in Kindler syndrome. Human molecular genetics 12, 925–935 (2003).

24 Lanschuetzer, C. M. et al. Characteristic immunohistochemical and ultrastructural findings indicate that Kindler’s syndrome is an apoptotic skin disorder. Journal of cutaneous pathology 30, 553–560 (2003).

25 Siegel, D. H. et al. Loss of kindlin-1, a human homolog of the Caenorhabditis elegans actin-extracellular-matrix linker protein UNC-112, causes Kindler syndrome. American journal of human genetics 73, 174–187 (2003).

26 Rognoni, E., Ruppert, R. & Fassler, R. The kindlin family: functions, signaling properties and implications for human disease. J Cell Sci 129, 17–27 (2016).

27 Guo, L. et al. Kindlin-2 links mechano-environment to proline synthesis and tumor growth. Nature communications 10, 845 (2019).

28 Mackinnon, A. C., Qadota, H., Norman, K. R., Moerman, D. G. & Williams, B. D. C. elegans PAT-4/ILK functions as an adaptor protein within integrin adhesion complexes. Current biology : CB 12, 787–797 (2002).

29 Theodosiou, M. et al. Kindlin-2 cooperates with talin to activate integrins and induces cell spreading by directly binding paxillin. eLife 5, e10130 (2016).

30 Kadry, Y. A., Huet-Calderwood, C., Simon, B. & Calderwood, D. A. Kindlin-2 interacts with a highly conserved surface of ILK to regulate focal adhesion localization and cell spreading. J Cell Sci 131 (2018).

31 Moser, M., Legate, K. R., Zent, R. & Fassler, R. The tail of integrins, talin, and kindlins. Science 324, 895–899 (2009).

32 Li, H. et al. Structural basis of kindlin-mediated integrin recognition and activation. Proceedings of the National Academy of Sciences of the United States of America 114, 9349–9354 (2017).

33 Harbury, P. B., Zhang, T., Kim, P. S. & Alber, T. A Switch between 2-Stranded, 3-Stranded and 4-Stranded Coiled Coils in Gcn4 Leucine-Zipper Mutants. Science 262, 1401–1407 (1993).

34 Li, P. et al. Phase transitions in the assembly of multivalent signalling proteins. Nature 483, 336–340 (2012).

35 Zeng, M. et al. Phase Transition in Postsynaptic Densities Underlies Formation of Synaptic Complexes and Synaptic Plasticity. Cell 166, 1163–1175 e1112 (2016).

36 Banjade, S. & Rosen, M. K. Phase transitions of multivalent proteins can promote clustering of membrane receptors. eLife 3 (2014).

37 Zeng, M. et al. Reconstituted Postsynaptic Density as a Molecular Platform for Understanding Synapse Formation and Plasticity. Cell 174, 1172–1187 e1116 (2018).

38 Mitrea, D. M. & Kriwacki, R. W. Phase separation in biology; functional organization of a higher order. Cell communication and signaling : CCS 14, 1 (2016).

39 Banani, S. F., Lee, H. O., Hyman, A. A. & Rosen, M. K. Biomolecular condensates: organizers of cellular biochemistry. Nature reviews. Molecular cell biology 18, 285–298 (2017).

40 Case, L. B., Ditlev, J. A. & Rosen, M. K. Regulation of Transmembrane Signaling by Phase Separation. Annual review of biophysics 48, 465–494 (2019).

41 Calderwood, D. A. et al. The Talin head domain binds to integrin beta subunit cytoplasmic tails and regulates integrin activation. The Journal of biological chemistry 274, 28071–28074 (1999).

42 Fukuda, K. et al. Molecular basis of kindlin-2 binding to integrin-linked kinase pseudokinase for regulating cell adhesion. The Journal of biological chemistry 289, 28363–28375 (2014).

43 Fukuda, K., Gupta, S., Chen, K., Wu, C. & Qin, J. The pseudoactive site of ILK is essential for its binding to alpha-Parvin and localization to focal adhesions. Molecular cell 36, 819–830 (2009).

44 Stutchbury, B., Atherton, P., Tsang, R., Wang, D. Y. & Ballestrem, C. Distinct focal adhesion protein modules control different aspects of mechanotransduction. J Cell Sci 130, 1612–1624 (2017).

45 Bouaouina, M. et al. A conserved lipid-binding loop in the kindlin FERM F1 domain is required for kindlin-mediated alphaIIbbeta3 integrin coactivation. The Journal of biological chemistry 287, 6979–6990 (2012).

46 Liu, Y., Zhu, Y., Ye, S. & Zhang, R. Crystal structure of kindlin-2 PH domain reveals a conformational transition for its membrane anchoring and regulation of integrin activation. Protein & cell 3, 434–440 (2012).

47 Yates, L. A. et al. Structural and functional characterization of the kindlin-1 pleckstrin homology domain. The Journal of biological chemistry 287, 43246–43261 (2012).

48 Ni, T. et al. Structure and lipid binding properties of the kindlin-3 pleckstrin homology domain. The Biochemical journal (2016).

49 Brangwynne, C. P. et al. Germline P granules are liquid droplets that localize by controlled dissolution/condensation. Science 324, 1729–1732 (2009).

50 Berry, J., Brangwynne, C. P. & Haataja, M. Physical principles of intracellular organization via active and passive phase transitions. Reports on progress in physics. Physical Society 81, 046601 (2018).

51 Su, X. et al. Phase separation of signaling molecules promotes T cell receptor signal transduction. Science 352, 595–599 (2016).

52 Ditlev, J. A. et al. A composition-dependent molecular clutch between T cell signaling condensates and actin. eLife 8 (2019).

53 Wu, X. et al. RIM and RIM-BP Form Presynaptic Active-Zone-like Condensates via Phase Separation. Molecular cell 73, 971–984 e975 (2019).

54 Beutel, O., Maraspini, R., Pombo-Garcia, K., Martin-Lemaitre, C. & Honigmann, A. Phase Separation of Zonula Occludens Proteins Drives Formation of Tight Junctions. Cell 179, 923–936 e911 (2019).

55 Schwayer, C. et al. Mechanosensation of Tight Junctions Depends on ZO-1 Phase Separation and Flow. Cell 179, 937–952 e918 (2019).

56 Case, L. B., Zhang, X., Ditlev, J. A. & Rosen, M. K. Stoichiometry controls activity of phase-separated clusters of actin signaling proteins. Science 363, 1093–1097 (2019).

57 Hato, T., Pampori, N. & Shattil, S. J. Complementary roles for receptor clustering and conformational change in the adhesive and signaling functions of integrin alphaIIb beta3. The Journal of cell biology 141, 1685–1695 (1998).

58 Wiseman, P. W. et al. Spatial mapping of integrin interactions and dynamics during cell migration by image correlation microscopy. J Cell Sci 117, 5521–5534 (2004).

59 Hart, R., Stanley, P., Chakravarty, P. & Hogg, N. The kindlin 3 pleckstrin homology domain has an essential role in lymphocyte function-associated antigen 1 (LFA-1) integrin-mediated B cell adhesion and migration. The Journal of biological chemistry 288, 14852–14862 (2013).

60 Bottcher, R. T. et al. Kindlin-2 recruits paxillin and Arp2/3 to promote membrane protrusions during initial cell spreading. The Journal of cell biology 216, 3785–3798 (2017).

61 Zhu, L. et al. Structural Basis of Paxillin Recruitment by Kindlin-2 in Regulating Cell Adhesion. Structure 27, 1686–1697 e1685 (2019).

62 Qadota, H., Moerman, D. G. & Benian, G. M. A molecular mechanism for the requirement of PAT-4 (integrin-linked kinase (ILK)) for the localization of UNC-112 (Kindlin) to integrin adhesion sites. The Journal of biological chemistry 287, 28537–28551 (2012).

63 Huet-Calderwood, C. et al. Differences in binding to the ILK complex determines kindlin isoform adhesion localization and integrin activation. J Cell Sci 127, 4308–4321 (2014).

64 Tantivejkul, K., Vucenik, I. & Shamsuddin, A. M. Inositol hexaphosphate (IP6) inhibits key events of cancer metastasis: II. Effects on integrins and focal adhesions. Anticancer research 23, 3681–3689 (2003).

65 Fu, C. et al. Neuronal migration is mediated by inositol hexakisphosphate kinase 1 via alpha-actinin and focal adhesion kinase. Proceedings of the National Academy of Sciences of the United States of America 114, 2036–2041 (2017).

66 Rojas, T. et al. Inositol hexakisphosphate kinase 3 promotes focal adhesion turnover via interactions with dynein intermediate chain 2. Proceedings of the National Academy of Sciences of the United States of America 116, 3278–3287 (2019).

67 Wang, Q.-S. et al. Upgrade of macromolecular crystallography beamline BL17U1 at SSRF. Nucl. Sci. Tech. 29, 68 (2018).

68 Otwinowski, Z. & Minor, W. Processing of X-ray Diffraction Data Collected in Oscillation Mode Methods in Enzymology 276, 307–326 (1997).

69 Adams, P. D. et al. PHENIX: a comprehensive Python-based system for macromolecular structure solution. Acta Crystallogr D Biol Crystallogr 66, 213–221 (2010).

70 Emsley, P. & Cowtan, K. Coot: model-building tools for molecular graphics. Acta Crystallogr D Biol Crystallogr 60, 2126–2132 (2004).

71 Davis, I. W. et al. MolProbity: all-atom contacts and structure validation for proteins and nucleic acids. Nucleic Acids Res 35, W375–383 (2007).

